# Suppression of ITPK1 and IPMK activities impairs mTORC1 signaling due to accelerated signal termination and prevents pancreatic ß-cell metabolic adaptation to excess glucose

**DOI:** 10.64898/2026.03.04.709646

**Authors:** Clarisse Iradukunda, Mary Virginia Whatley, Edward A. Salter, Dilipkumar Uredi, Xiaodong Wang, Joel F. Andrews, Andrzej Wierzbicki, Lucia E. Rameh

## Abstract

mTORC1 integrates growth factor and nutrient signals to regulate cellular metabolism, yet there are no metabolites known to directly regulate mTORC1 activity in cells. Cryo-EM studies revealed that inositol hexakisphosphate (IP_6_) associates with the FAT domain of mTOR, suggesting that inositol phosphates may directly modulate mTOR activity. We previously showed that higher-order inositol phosphates enhance mTORC1 kinase activity and stability *in vitro*. Here, we investigated whether inositol phosphate metabolism regulates mTORC1 signaling in HEK293 and pancreatic β-cells. Suppression or acute inhibition of inositol phosphate multikinase (IPMK) selectively reduced cellular IP_5_ levels without altering IP_6_ and resulted in reduced basal and insulin-stimulated mTORC1 signaling, without decreasing mTORC1 complex stability or lysosomal localization. ITPK1 knockdown phenocopied IPMK inhibition by reducing IP_5_ and basal mTORC1. Combined inhibition of IPMK and ITPK1 nearly abolished IP_5_ and reduced IP_6_, demonstrating that these enzymes compensate to supply IP_5_ for IP_6_ synthesis. Importantly, depletion of IP_5_ did not impair PI3K/Akt activation but accelerated termination of the mTORC1 signal, indicating a role for IP_5_ in stabilizing the active mTORC1 complex. Reduction of IP_5_ levels did not prevent insulin- or glucose-induced mTORC1 activation, revealing that inositol phosphates primarily regulate signal persistence rather than initiation. Lastly, we show that suppression of IPMK or ITPK1 in ß-cells delayed the mTORC1-dependent metabolic acceleration in response to excess glucose. Together, these findings identify higher order inositol phosphates as metabolic regulators that prolong mTORC1 activity, providing a potential strategy for modulating mTORC1 signaling in cells.

**Significance Statement:** mTORC1 is a central metabolic regulator whose chronic activation contributes to metabolic disease, yet mechanisms that sustain mTORC1 activity after its activation are poorly understood. We show that enzymes controlling inositol phosphate metabolism regulate the persistence of mTORC1 signaling in pancreatic β-cells by maintaining cellular levels of inositol pentakisphosphate (IP_5_). Reducing IP_5_ impairs basal and sustained mTORC1 signaling without affecting upstream growth factor or energy-sensing pathways, revealing a mechanism that controls signal duration rather than activation. Reduction in IP_5_ levels also delayed the mTORC1-dependent ß-cell metabolic acceleration in response to elevated glucose. These findings identify IP_5_ as a metabolic regulator of mTORC1 and suggest that targeting inositol phosphate metabolism may provide a strategy to modulate mTORC1 activity in metabolic disease.

## Introduction

mTORC1 is a central regulator of cellular metabolism, coordinating anabolic growth with nutrient and growth factor availability (1). Canonical activation of mTORC1 requires growth factor–dependent activation of the small GTPase RHEB and amino acid–dependent recruitment of the complex to lysosomal membranes. While these mechanisms explain how mTORC1 is acutely activated, they do not fully account for how mTORC1 activity is sustained beyond the lysosomal surface to reach other subcellular locations where its substrates reside (1). Additionally, unconventional pathways for mTORC1 hyperactivation by glucose have been described but the metabolites directly involved have not been identified (2–4).

Recent studies have suggested that inositol phosphates, which are small metabolites derived from dietary inositol or glucose, may directly modulate mTOR activity (5). Cryo-electron microscopy analyses revealed that inositol hexakisphosphate (IP_6_) occupies a positively charged pocket within the FAT domain of mTOR, named the I-site, and mutations disrupting this pocket impair kinase activity and/or protein stability (6, 7). These findings raised the possibility that IP_6_ functions either as structural cofactor for mTOR or as regulator of its activity. Consistent with the latter, we recently demonstrated that higher-order inositol phosphates enhance the activity, stability, and turnover of purified mTOR and mTORC1 *in vitro*, in a dose-dependent and reversible manner, with IP_5_ and IP_6_ being the most potent activators (5). However, whether inositol phosphate metabolism regulates mTORC1 signaling in cells, and which specific inositol phosphate species is/are functionally relevant, has not been addressed.

In cells, IP_6_ is synthesized through sequential phosphorylation of lower-order inositol phosphates, with IP_5_ serving as the immediate precursor (8). Generation of IP_5_ can occur through two distinct enzymatic pathways catalyzed by inositol phosphate multikinase (IPMK) and inositol trisphosphate kinase 1 (ITPK1) (9). IPMK has previously been implicated in mTORC1 regulation through kinase-independent scaffolding functions and through modulation of upstream signaling pathways, complicating efforts to assess the role of its inositol phosphate products (10). Moreover, the contribution of ITPK1 to mTORC1 signaling has not been examined. Based on our *in vitro* data, we hypothesized that IP_5_ generated by either IPMK or ITPK1 stabilizes active mTOR to promote sustained mTORC1 signaling in cells.

Pancreatic β-cells provide a physiologically relevant system in which to address this question. In these cells, mTORC1 activity is tightly linked to nutrient availability and insulin demand (11), and chronic elevation of basal mTORC1 signaling contributes to β-cell metabolic adaptation to hyperglycemia (2) and eventually to ß-cell dysfunction and failure (4, 12). We previously showed that chronic exposure to excess glucose induces mTORC1 hyperactivation in β-cells whereas acute glucose exposure dose-dependently induces mTORC1 through a mechanism that does not involve further stimulation of PI3K/Akt or suppression of AMPK, suggesting the existence of non-canonical regulatory inputs (2).

Here, we show that inositol phosphate metabolism regulates mTORC1 signaling in the pancreatic β-cell line INS-1, primary islet cells and HEK293 cells. Suppression or acute inhibition of IPMK, as well as knockdown of ITPK1, reduced cellular IP_5_ levels and impaired basal and growth factor-stimulated mTORC1 signaling without affecting PI3K/Akt activation, mTOR lysosomal recruitment or mTORC1 stability. IPMK and ITPK1 compensated for one another to maintain IP_5_ and IP_6_ synthesis, and combined inhibition of both pathways further impaired mTORC1 signal, especially under basal conditions (i.e. physiological glucose and low insulin). Although, acute glucose-induced mTORC1 activation was preserved and AMPK status was unchanged, the long-term metabolic adaptation to elevated glucose was impaired in cells with ITPK1 knockdown and/or IPMK inhibition. Interestingly, IPMK inhibition accelerated the de-activation of insulin-induced mTORC1 signal and the normalization of basal mTORC1 upon excess glucose removal. These findings identify IP_5_ as a metabolic regulator that prolongs mTORC1 signaling in β-cells.

## Results

### 1) Suppression of IPMK expression or inhibition of its catalytic activity reduces IP_5_ and impairs mTORC1 signaling in pancreatic ß-cells

To determine whether IPMK-dependent inositol phosphate synthesis contributes to mTORC1 signaling in pancreatic ß-cells, we knocked down IPMK expression in INS-1 cells using shRNA and measured cellular inositol phosphate levels and mTORC1 signaling. Partial knockdown of IPMK (Fig S1A) resulted in a significant reduction in IP_5_ levels and a smaller decrease in IP_4_, while IP_6_ and IP_7_ levels were unchanged (Fig. 1A and Fig. S1 B-C). Although total IP_4_ reduction narrowly missed statistical significance, analysis of IP_4_ subpeaks revealed a significant decrease in one IP_4_ species (Fig. S1 D-F).

**Figure 1.**
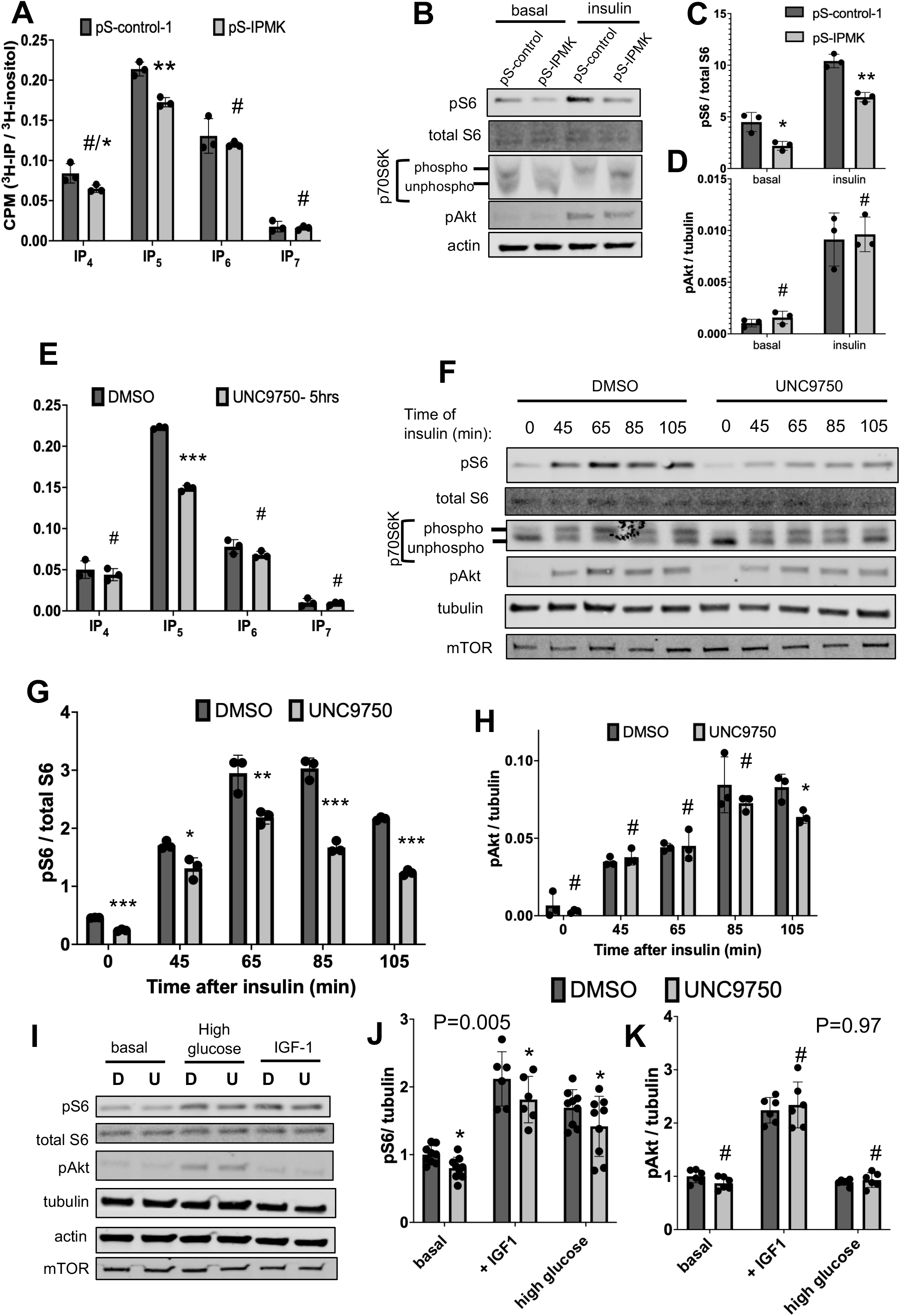
Knockdown or inhibition of IPMK catalytic activity in pancreatic ß-cells reduced IP_5_ and impaired mTORC1 signaling without affecting IP_6_ or Akt activation. **(A)** Cellular [³H]-labeled inositol phosphate levels in control (pS-control) or IPMK knockdown (pS-IPMK) INS-1 cells, normalized to [³H]-inositol. IPMK knockdown reduced IP_5_ and IP_4_, with no significant change in IP_6_ or IP_7_. **(B)** Representative immunoblots showing basal and insulin-stimulated mTORC1 signaling in control or IPMK knockdown cells, assessed by S6 phosphorylation and p70S6K mobility shift, and upstream insulin signaling assessed by Akt phosphorylation. **(C)** Quantification of S6 phosphorylation from three biologically independent samples, normalized to total S6. **(D)** Quantification of Akt phosphorylation from the same experiments, normalized to tubulin. Data are mean ± SD. Statistical significance is indicated in panels; (*)P=0.01-0.05; (**)P=0.001-0.002; (***) P<0.0003; (*/#) P=0.057; (#) P>0.06. **(E)** Cellular [³H]-labeled inositol phosphate levels in INS-1 cells treated with vehicle control (DMSO) or UNC9750 for 5 hrs and normalized to [³H]-inositol. IPMK inhibition reduced IP_5_ and IP_4_, with no significant change in IP_6_ or IP_7_. **(F)** Representative immunoblots showing basal and time dependent insulin-stimulated mTORC1 signaling, assessed by S6 phosphorylation and p70S6K mobility shift, and upstream insulin signaling assessed by Akt phosphorylation. Also shown are the levels of mTOR. **(G)** Quantification of S6 phosphorylation from three biologically independent samples, normalized to total S6. **(H)** Quantification of Akt phosphorylation from the same experiments, normalized to tubulin. **(I)** Representative immunoblots showing basal, glucose-stimulated and IGF-1-stimulated mTORC1 signaling in human islets treated with vehicle (D) or UNC9750 (U), assessed by S6 phosphorylation and upstream IGF-1 signaling assessed by Akt phosphorylation. Also shown are the levels of mTOR. Data are mean ± SD. Statistical significance is indicated in panels. (*)P=0.02-0.04; (**)P=0.016; (***) P<0.0003; (#) P>0.05. **(J)** Quantification of S6 phosphorylation from three biologically independent samples obtained from three separate donors, and normalized to tubulin. **(K)** Quantification of Akt phosphorylation from the same experiments, normalized to tubulin. Data are mean ± SD. Statistical significance is indicated in panels. (*)P=0.017-0.035; (#) P>0.08. Two-way ANOVA shows P=0.005 for pS6 measurements (D versus U) and P=0.97 for pAkt measurements.

We next assessed mTORC1 signaling in INS-1 cells preconditioned in physiological glucose. IPMK knockdown reduced both basal and insulin-stimulated mTORC1 signaling, as measured by S6 phosphorylation and p70S6K mobility shift (Fig. 1 B-C). In contrast, basal and insulin-stimulated Akt phosphorylation were unchanged (Fig. 1B and D), indicating intact upstream PI3K signaling. Similar inhibition of mTORC1 signaling was observed following transient IPMK knockdown using siRNA (Fig. S2 A-D), confirming that the effects were not due to long-term alterations in cell proliferation.

To determine whether IPMK catalytic activity is required for its effect on mTORC1 signaling, we treated INS-1 cells with the selective IPMK inhibitor UNC9750 (13). Extended treatment during [^3^H]-inositol metabolic labeling reduced IP_5_ by ∼80% and IP_4_ by ∼16%, without altering IP_6_ or IP_7_ levels (Fig. S3 A-C). Short treatment reduced pre-labeled IP_5_ by ∼33% with no significant change in other inositol phosphates (Fig. 1E and S3D). Under these conditions, basal and insulin-stimulated mTORC1 signaling were significantly reduced (Fig. 1F and G), whereas early Akt phosphorylation was preserved (Fig. 1F and H). mTORC1 activation kinetics in response to insulin were unchanged, indicating that IPMK inhibition affects signal magnitude rather than activation timing. Notably, IPMK inhibition caused a modest but statistically significant decrease in Akt phosphorylation after prolonged insulin stimulation, which indicate partial effect on sustained mTORC2 activation.

To confirm the effect of IPMK inhibition on mTORC1 signaling in pancreatic islet cells, we treated dissociated human islets with UNC9750 for 3 hrs. As for INS-1 cells, mTORC1 signaling (basal, growth factor- or glucose-stimulated) was significantly reduced by the inhibitor (Fig. 1 I-J), whereas early Akt phosphorylation was preserved (Fig. 1I and K).

Together, these results show that suppression of IPMK activity selectively reduces IP_5_ and impairs basal and growth factor-stimulated mTORC1 signaling in both cultured ß-cells and pancreatic islets, independently of upstream growth factor signaling.

### 2) IPMK inhibition reduces mTORC1 without affecting mTOR stability or mTOR/Raptor interaction

Mutant mTOR in which 5 of the positively charged residues that coordinate IP_6_ were replaced by negatively charged residues failed to express a stable protein, suggesting that inositol phosphate binding stabilizes mTOR (6). Yet, mutant mTOR with 2 or 3 substitutions in the I-site expressed active kinase in the presence of mLST8, SIN and Rictor (6), indicating that IP_6_ binding is dispensable for proper folding in the context of the full complex. In our studies, IPMK inhibition did not change mTOR expression (Fig 1 F and I) suggesting that reduced mTORC1 signaling cannot be explained by decreased mTOR stability in cells. To address whether mTOR association with the regulatory subunit Raptor was affected by IPMK inhibition, we immunoprecipitated mTOR from cells treated with DMSO or UNC9750. UNC9750 did not affect the ability of mTOR to form a complex with Raptor confirming that stability of mTORC1 is not affected by IPMK inhibition (Fig S4 A-B).

### 3) IPMK inhibition reduces IP_5_ and mTORC1 signaling in HEK293 cells without affecting lysosomal recruitment

Next, we examine whether IPMK inhibition can affect mTORC1 signaling in other cell types. Treatment of [^3^H]-inositol-labeled HEK293 cells with UNC9750 for 5 hrs reduced IP_5_ levels by 43% without significant changes in IP_4_, IP_6_ or IP_7_ (Fig.2 A). Basal mTORC1 signaling was inhibited by 33-78% in treated cells, as judged by p70S6K mobility shift or S6 phosphorylation, respectively (Fig 2B-D). IGF-1-induced mTORC1 signaling was also significantly reduced (40-44%). There was no effect of UNC9750 on basal pAkt levels, whereas IGF-1-induced phospho-Akt had a 21% reduction (Fig. 2B and E), which could be due to either inhibition of mTORC2 or a reduction in PIP_3_ (phosphatidylinositol-3,4,5-P_3_) synthesis (14).

**Figure 2:**
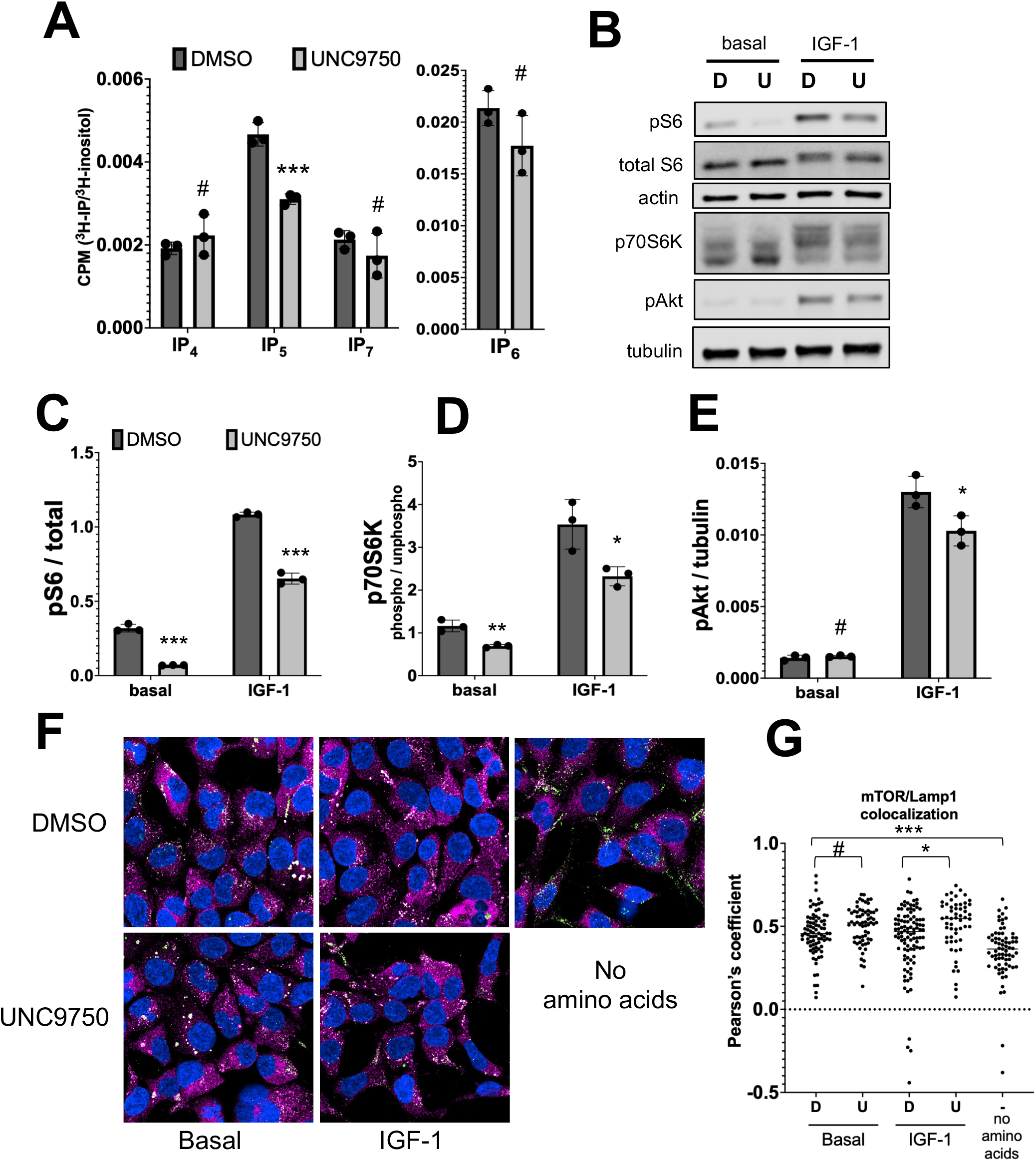
IPMK inhibition in HEK293 cells reduced IP_5_ levels and suppressed mTORC1 signaling without impairing mTOR co-localization with lysosomes. **(A)** Cellular [³H]-labeled inositol phosphate levels in DMSO- or UNC9750-treated HEK293 cells, normalized to [³H]-inositol. IPMK inhibition reduced IP_5_, with no significant change in IP_4_, IP_6_ or IP_7_. **(B)** Representative immunoblots showing basal and IGF-1-stimulated mTORC1 signaling in cells treated with DMSO (D) or UNC9750 (U), assessed by S6 phosphorylation and p70S6K mobility shift, and upstream insulin signaling assessed by Akt phosphorylation. **(C)** Quantification of S6 phosphorylation from three biologically independent samples, normalized to total S6. **(D)** Quantification of p70S6K phosphorylation (top bands/lower band) from the same experiment as in C. **(E)** Quantification of Akt phosphorylation from the same experiment as in C, normalized to tubulin. Data are mean ± SD. Statistical significance is indicated in panels. (*)P=0.02-0.04; (**)P=0.004; (***) P<0.0009; (#) P>0.05. **(F)** Immunofluorescence images of HEK293 cells treated with DMSO or UNC9750 and stimulated or not with IGF-1. mTOR staining is shown in magenta and Lamp1 in green. Co-localization is indicated in white. Also shown are control cells starved for amino acids for 30 minutes. **(G)** Cell-based quantification of mTOR/Lamp1 co-localization from the same experiments shown in F, as measured by Pearson’s coefficient. One-way ANOVA P=0.0001, Turkey’s multiple comparison test (*)P=0.023; (***)P=0.0004; (#) P=0.39.

The robust reduction in basal mTORC1 signaling by IPMK inhibition without an effect on basal Akt phosphorylation is consistent with a direct role for IP_5_ in mTORC1 activation. To determine whether IPMK inhibition affects the subcellular localization of mTOR to the lysosomal region where mTORC1 activation by RHEB takes place (1), we examined the effect of UNC9750 on mTOR colocalization with the lysosomal marker Lamp1. UNC9750 treatment did not decrease mTOR colocalization with Lamp1 (Fig.2 F-G and S4 C-D), confirming that the effect of IPMK inhibition on mTORC1 signaling cannot be explained by impaired recruitment of mTOR to the lysosome. Instead, there was a small increase in mTOR/Lamp1 co-localization upon IPMK inhibition (Fig. 2G) which is consistent with previous findings that mTORC1 activity is necessary for its release from the lysosomal surface (15).

### 4) Suppression of ITPK1 expression reduces IP_5_ and impairs mTORC1 signaling in pancreatic ß-cells

To determine whether reduced mTORC1 signaling reflects loss of inositol phosphates rather than IPMK-specific functions, we suppressed ITPK1 as an independent strategy to reduce IP_5_. Knockdown of ITPK1 in INS-1 cells reduced IP_4_ and IP_5_ levels by approximately 30% and 47%, respectively, without affecting IP_6_ or IP_7_ (Fig. 3A and Fig. S5 A-C).

**Figure 3:**
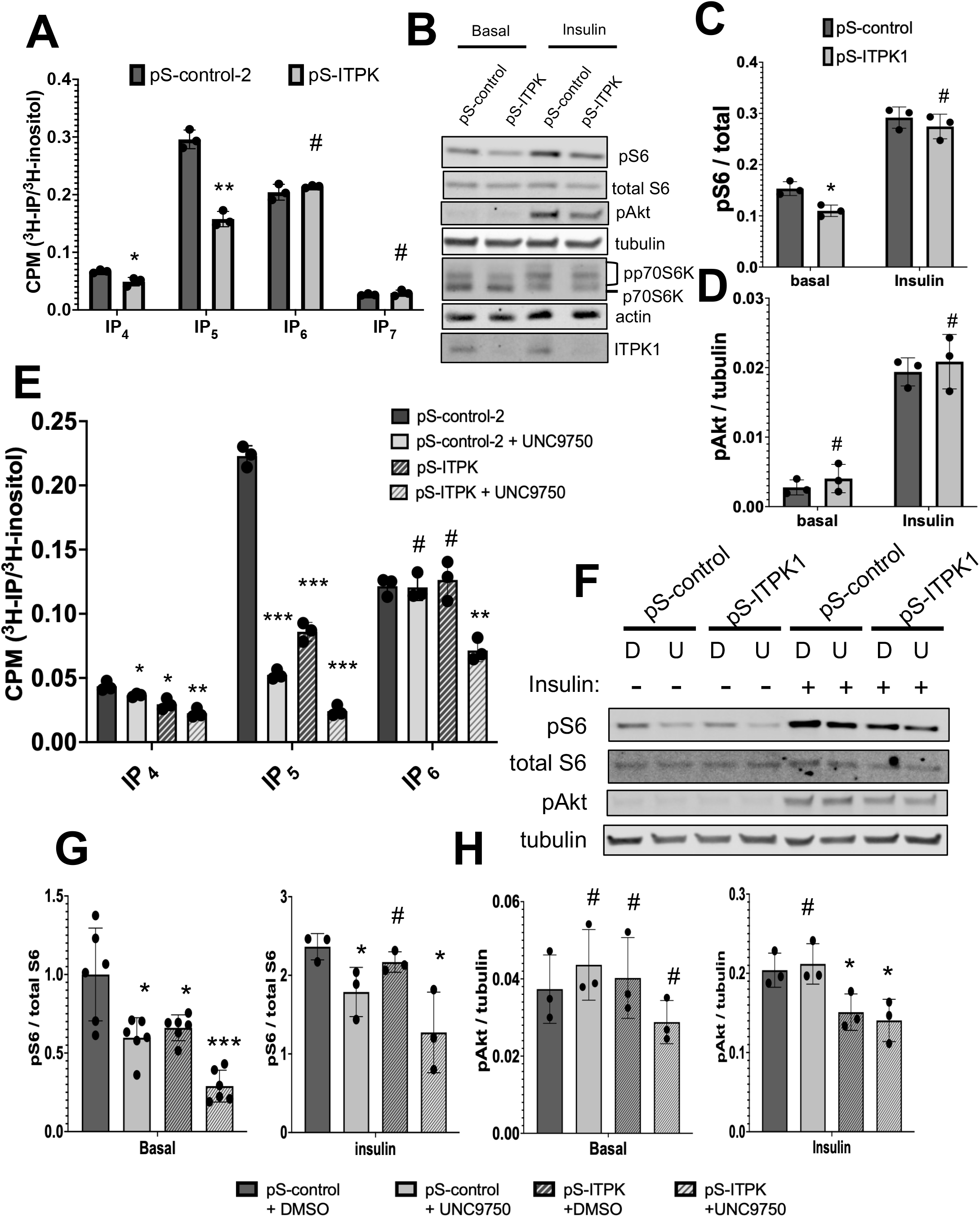
ITPK1 knockdown reduced IP_4_ and IP_5_ levels and suppressed mTORC1 signaling. **(A)** Cellular [³H]-labeled inositol phosphate levels in control (pS-control-2) or ITPK1 knockdown (pS-ITPK) INS-1 cells, normalized to [³H]-inositol. ITPK1 knockdown reduced IP_5_ and IP_4_, with no significant change in IP_6_ or IP_7_. **(B)** Representative immunoblots showing basal and insulin-stimulated mTORC1 signaling in control or ITPK1 knockdown cells, assessed by S6 phosphorylation, p70S6K mobility shift, and upstream insulin signaling assessed by Akt phosphorylation. Also shown are the levels of ITPK1. **(C)** Quantification of S6 phosphorylation from three biologically independent samples, normalized to total S6. **(D)** Quantification of Akt phosphorylation from the same samples as in C, normalized to tubulin. **(E)** Cellular [³H]-labeled inositol phosphate levels in control (pS-control) or ITPK1 knockdown (pS-ITPK1) INS-1 cells treated with vehicle control (DMSO) or UNC9750, normalized to [³H]-inositol. IPMK inhibition with ITPK1 knockdown had additive effects in reducing IP_4_ and IP_5_, and synergistic effect in reducing IP_6_. **(F)** Representative immunoblots showing basal and insulin-stimulated mTORC1 signaling in control (pS-control) or ITPK1 knockdown (pS-ITPK1) cells treated with DMSO (D) or UNC9750 (U), assessed by S6 phosphorylation, and upstream insulin signaling assessed by Akt phosphorylation **(G)** Basal and insulin-stimulated mTORC1 signaling, assessed by S6 phosphorylation. Quantification of S6 phosphorylation from three or six biologically independent samples, normalized to total S6. **(H)** Basal and insulin-stimulated Akt activation as a measure of signaling upstream of mTORC1. Quantification of Akt phosphorylation from the same experiments, normalized to tubulin. Data are mean ± SD. Statistical significance is indicated in panels. (*)P=0.01-0.05; (**)P=0.001-0.002; (***) P<0.0002; (#) P>0.05.

Knockdown of ITPK1 protein (Fig. 3B and S5E) significantly reduced basal, but not insulin-induced mTORC1 signaling (Fig. 3B-C, 3F-G and S5D). Akt phosphorylation remained unchanged under basal conditions (Fig. 3B, D, F and H). Insulin-dependent Akt phosphorylation was either unchanged (Fig 3D) or slightly blunted after longer insulin stimulation of ITPK1 knockdown cells (Fig 3F and H). Thus, ITPK1 knockdown phenocopied IPMK knockdown/inhibition. These data demonstrate that depletion of IP_5_ via a pathway independent of IPMK is sufficient to impair mTORC1 signaling and perhaps mTORC2.

### 5) ITPK1 and IPMK compensate for each other to maintain IP_5_ levels and mTORC1 signaling

Because IP_6_ levels were preserved despite substantial reductions in IP_5_ following either IPMK inhibition or ITPK1 knockdown, we tested whether IPMK and ITPK1 compensate to maintain inositol phosphate synthesis. Inhibition of IPMK in ITPK1 knockdown cells further reduced IP_5_ levels from 61% to 89% and resulted in a significant decrease in IP_6_ (Fig. 3E and S6A-B). In contrast, ITPK1 knockdown or IPMK inhibition alone had no effect on IP_6_ levels. IP_4_ levels were reduced additively by combined ITPK1 knockdown and IPMK inhibition (Fig.3E), whereas IP_3_ and IP_7_ levels remained unchanged (Fig. S6C). These results indicate that in INS-1 cells IPMK and ITPK1 have redundant roles in supplying IP_5_ for IP_6_ synthesis and that neither pathway alone is rate limiting.

### 6) Reduction in IP_5_ correlates with impaired mTORC1 signaling

We next examined how combined depletion of IPMK- and ITPK1-derived inositol phosphates affects mTORC1 signaling in INS-1 cells. In ITPK1 knockdown cells treated with UNC9750, basal and insulin-stimulated mTORC1 signaling were reduced by approximately 80% and 50%, respectively (Fig. 3 F and G). Akt phosphorylation was modestly reduced following prolonged insulin stimulation but remained unchanged under basal conditions (Fig. 3H), suggesting limited effects on mTORC2 relative to mTORC1. Across all perturbations, the degree of mTORC1 inhibition closely correlated with the extent of IP_5_ depletion, supporting a functional relationship between IP_5_ levels and mTORC1 signaling.

### 7) IPMK inhibition accelerates the decline in mTORC1 signaling

We next asked whether reduced inositol phosphate levels affect the persistence of mTORC1 signaling after termination of upstream PI3K activity. Cells were stimulated with insulin and subsequently treated with either wortmannin to inhibit mTORC1 activation at the level of PI3K, or rapamycin to directly terminate the mTORC1 signal. The rate of signal decline was measured over time. Wortmannin rapidly abolished Akt phosphorylation within 20 minutes, and this was unaffected by IPMK inhibition (Fig. 4A and B), confirming that the upstream signal was rapidly terminated. mTORC1 signaling declined more gradually, requiring up to 60 minutes for substantial loss of phosphorylated p70S6K and S6 in control cells (Fig. 4A, C and D). Notably, IPMK inhibition accelerated mTORC1 signal decay with an abrupt loss of signal at 20 minutes post-wortmannin, as compared to control.

**Figure 4:**
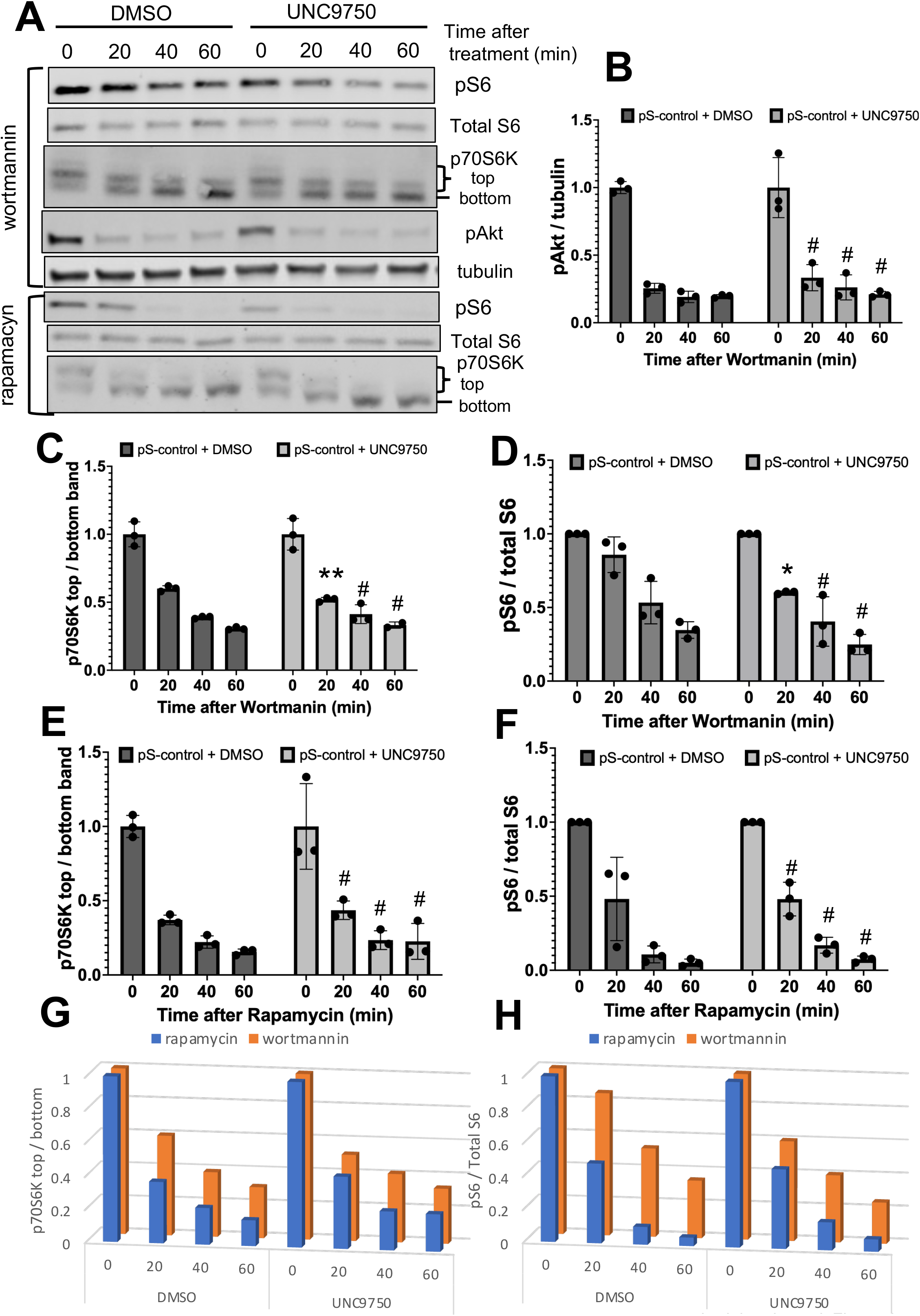
IPMK inhibition accelerates the decay of mTORC1 signaling. **(A)** Representative western blots showing the decline in mTORC1 signaling after signal termination with wortmannin or rapamycin in INS-1 cells treated with DMSO or UNC9750, assessed by p70S6K mobility shift or S6 phosphorylation (pS6), and upstream insulin signaling assessed by Akt phosphorylation (pAkt). **(B)** Quantification of Akt phosphorylation from three biologically independent samples, normalized to tubulin. Shown are the rate of signal decay after **wortmannin** treatment normalized by time 0 for each treatment group. **(C)** Quantification of p70S6K phosphorylation from three biologically independent samples, as a ratio between upper phosphorylated bands over bottom, unphosphorylated band. Shown are the rate of signal decay after **wortmannin** treatment normalized by time 0 for each treatment group. **(D)** Quantification of S6 phosphorylation from three biologically independent samples, normalized to total S6. Shown are the rate of signal decay after **wortmannin** treatment normalized by time 0 for each treatment group. **(E)** Quantification of p70S6K phosphorylation from three biologically independent samples, as a ratio between upper phosphorylated bands over bottom, unphosphorylated band. Shown are the rate of signal decay after **rapamycin** treatment normalized by time 0 for each treatment group. **(F)** Quantification of S6 phosphorylation from three biologically independent samples, normalized to total S6. Shown are the rate of signal decay after **rapamycin** treatment normalized by time 0 for each treatment group. **(G)** 3-D graph showing the juxtaposition of graphs C (wortmannin treatment in orange) and E (rapamycin treatment in blue). **(H)** 3-D graph showing the juxtaposition of graphs D (wortmannin treatment in orange) and F (rapamycin treatment in blue). Data are mean ± SD. Statistical significance is indicated in panels. (*)P=0.01-0.05; (**)P=0.005; (#) P>0.05.

mTORC1 signal decline is the result of both the deactivation of mTORC1 kinase and dephosphorylation of its downstream substrates. To determine whether IPMK inhibition affected the rate of substrate dephosphorylation, we measured the loss of phosphorylated p70S6K and S6 after direct inhibition of mTORC1 with rapamycin. Phosphorylated p70S6K was drastically reduced at 20 minutes post-rapamycin and this decline was not accelerated by IPMK inhibition (Fig 4 A and E). The rate of S6 dephosphorylation was slightly delayed as compared to p70S6K, reaching completion after 40 minutes of rapamycin, and IPMK inhibition had no effect on this kinetics (Fig 4 A and F). Thus, it is unlikely that reduced IP_5_ levels increase phosphatase activity.

The juxtaposition of the graphs showing the decay in mTORC1 signaling after rapamycin or wortmannin (Fig. 4G and H) highlights the persistence in mTORC1 signal after PI3K inhibition as compared to the expected decay in signal after rapamycin (difference between the orange and the blue bars). It also highlights that in cells treated with IPMK inhibitor, the decay in mTORC1 signal approximates the expected decay due to dephosphorylation. Similar results were observed in ITPK1 knockdown cells treated with IPMK inhibitor, whereas ITPK1 knockdown alone did not significantly alter signal decay kinetics (Fig. S7 A-H).

Together, these results indicated that active mTORC1 is stabilized after Akt signal termination and that this is dependent on IPMK kinase activity.

### 8) Reduction in IPMK/ITPK1-derived inositol phosphates impairs mTORC1-dependent chronic adaptation to elevate glucose

We have previously shown that sustained mTORC1 signaling is necessary for ß-cells to metabolically adapt to chronic elevated glucose (2). Thus, we examined whether IPMK- and/or ITPK1-dependent reduction in mTORC1 signaling affects the metabolic response of ß-cells to elevated glucose. INS-1 cells preconditioned in physiological glucose were stimulated with high glucose in the presence or absence of IPMK inhibition. Acute glucose stimulation did not alter Akt or AMPK phosphorylation, as previously reported (2), and neither did IPMK inhibition (Fig. S8A, E, and F).

Consistent with previous results, UNC9750 treatment reduced basal and glucose-stimulated mTORC1 signaling, although the fold induction in response to glucose was preserved (Fig. 5A and Fig. S8A-C). Similar results were obtained following IPMK or ITPK1 knockdown, as well as combined perturbation (Fig. 5B and C). Thus, reduced ITPK1 and IPMK activities dampen glucose-dependent mTORC1 signaling but are not required for glucose-induced activation.

**Figure 5:**
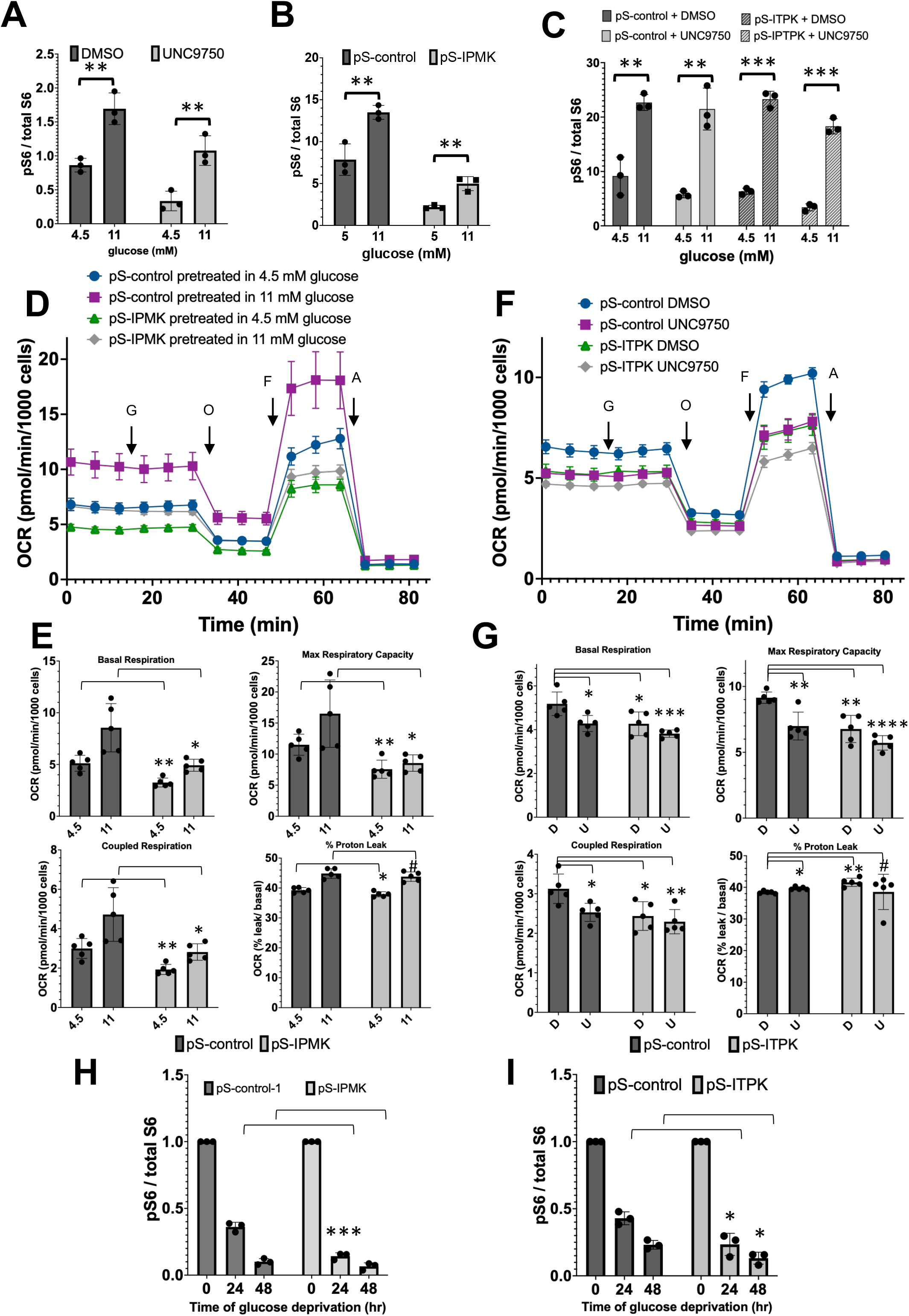
Suppression of IP_5_ impairs metabolic adaptation to chronic glucose and accelerates the decline in glucose-dependent basal mTORC1 activity, without affecting acute response to glucose. **(A-C)** Basal (4.5 mM) and glucose-stimulated (11 mM, 30 min) mTORC1 signaling, assessed by S6 phosphorylation, in INS-1 cells treated with DMSO or UNC9750 **(A and C)**, in control (pS-control) or IPMK knockdown (pS-IPMK) cells **(B)** and in control or ITPK1 knockdown (pS-ITPK1) cells **(C)**. Darta are the quantification of S6 phosphorylation from three biologically independent samples, normalized to total S6. **(D)** Representative experiment showing measurements of oxygen consumption rate (OCR) of control (pS-control) or IPMK knockdown (pS-IPMK) INS-1 cells cultured in media containing 4.5 or 11 mM glucose for 48 hrs before mitochondrial stress test. Starting measurements were performed in media containing 3 mM glucose followed by injections of 8 mM glucose (G), oligomycin B (O), FCCP (F) and antimycin A (A), as indicated by the arrows. Data are the mean and SEM of 5 independent biological samples. **(E)** Calculated values for basal respiration (at 3 mM glucose), maximal respiratory capacity, ATP-coupled respiration and percentage proton leak from the experiment shown in D. **(F)** Representative experiment showing measurements of oxygen consumption rate (OCR) of control (pS-control) or ITPK1 knockdown (pS-ITPK1) INS-1 cells cultured in media containing 11 mM glucose with DMSO or UNC9750 for 48 hrs before mitochondrial stress test. Starting measurements were performed in media containing 3 mM glucose followed by injections of 8 mM glucose (G), oligomycin B (O), FCCP (F) and antimycin A (A), as indicated by the arrows. Data are the mean and SEM of 5 independent biological samples. **(G)** Calculated values for basal respiration (at 3 mM glucose), maximal respiratory capacity, ATP-coupled respiration and percentage proton leak from the experiment shown in F. **(H)** Decline in basal mTORC1 signaling upon glucose withdrawal (from 11 mM to 4.5 mM) of control or IPMK knockdown INS-1 cells, as assessed by S6 phosphorylation. Quantification of S6 phosphorylation from three biologically independent samples, normalized to total S6. **(I)** Decline in basal mTORC1 signaling upon glucose withdrawal (from 11 mM to 4.5 mM) of control or ITPK1 knockdown INS-1 cells, as assessed by S6 phosphorylation. Quantification of S6 phosphorylation from three biologically independent samples, normalized to total S6. Data are mean ± SD. Statistical significance is indicated in panels. (*)P=0.01-0.05; (**)P=0.002-0.009; (***) P<0.0008; (#) P>0.05.

Next, we examined the effect of IPMK knockdown on glucose-induced metabolic acceleration of ß-cells, an mTORC1-dependent response (2). INS-1 cells with or without IPMK knockdown were preconditioned in physiological glucose and then chronically exposed to elevated glucose for 48 hrs, which is sufficient to increase basal respiration (at 3 mM glucose), coupled respiration, proton leak and max respiratory capacity (at 8 mM glucose) (Fig. 5D and E). IPMK knockdown cells had overall lower metabolic rate with decreased basal and coupled respiration and max respiratory capacity. Importantly, the response to chronically elevated glucose was impaired but not completely abolished. Cells with ITPK1 knockdown also had a lower metabolic rate after exposure to chronic elevated glucose as compared to control cells, with lower basal (at 3 mM glucose) and coupled respiration and max respiratory capacity (at 8 mM glucose) (Fig. 5F and G). IPMK inhibition with UNC9750 further decreased the metabolic rate of control INS-1 or ITPK1 knockdown cells (Fig. 5 F and G). These results demonstrate that IPMK and/or ITPK1 activities contribute to ß-cells long-term metabolic adaptation to chronic elevated glucose, most likely through inositol phosphate-dependent regulation of mTORC1.

We have previously shown that INS-1 cells chronically growing in excess glucose (11 mM) have high basal mTORC1 which is lowered after 48 hrs in physiological glucose (4-5 mM) (2). Thus, we addressed whether suppression of IPMK and/or ITPK1 accelerates the recovery of low basal mTORC1 levels upon removal of excess glucose. INS-1 cells pre-conditioned in 11 mM glucose were transferred to 4.5 mM glucose and the rate of recovery was measured. In cells with IPMK or ITPK1 knockdown, basal mTORC1 levels were completely reduced after 24 hrs in low glucose, whereas control cells required 48hrs (Fig. 5H and I). These results indicate that inositol phosphates derived from IPMK or ITPK1 prolong the active state of mTORC1 induced by chronic excess glucose.

### 9) IP_5_ binds the I-site of the mTOR FAT domain with strength comparable to that of the singly-protonated species of IP_6_, according to quantum-based modeling

Our results show that reduction in cellular IP_5_ levels closely correlate with reduced mTORC1 signal, even when IP_6_ levels are unchanged. We used quantum-based modeling to address putative IP_5_ binding modes to the I-site of mTOR. The cryo-EM structure of mTORC2:IP_6_ (pdb entry 6ZWO) shows that only 5 phosphate groups of IP_6_ make contact with positively-charged residues of the I-site; the sixth (on C4) is positioned at a gap (Fig. 6A). Thus, we explored orientations of IP_5_ within the I-site to see if its stereo arrangement of 5 phosphates can produce strong binding. In the most favored orientation of IP_5_ (Fig. 6B), the inositol scaffold is rotated vis-a-vis IP_6_, with its C2 hydroxyl group positioned at the gap. As summarized in Table 1, IP_5_ makes a similar number of phosphate/cation contacts within the I-site and binds more strongly than doubly-protonated IP_6_ (H_2_IP_6_) and with about the same strength as singly-protonated IP_6_ (HIP_6_). Fully-deprotonated IP_6_ is the strongest binding species, but only by ∼7.6% over IP_5_. For IP_4_, with the additional loss of the equatorial phosphate on C3, the binding energy drops by ∼12% compared to IP_5_.

**Figure 6:**
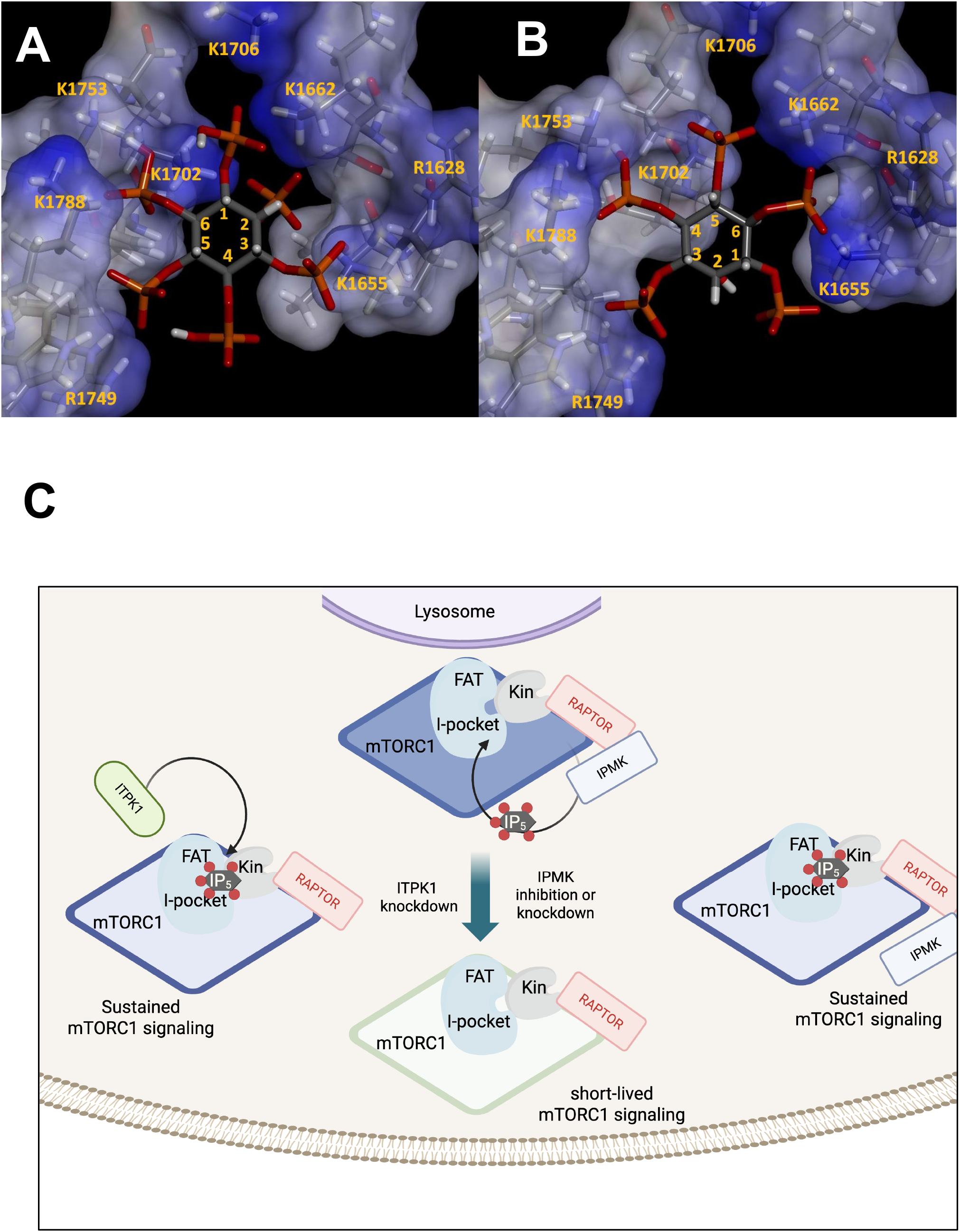
Predicted IP_5_ binding within the I-site of the mTOR FAT domain and role in mTORC1 signaling. **(A)** Binding of IP_6_ to the I-site of mTOR, according to (Scaiola et al., 2020). **(B)** Binding of IP_5_ to the I-site of mTOR, according to quantum-based modeling. The optimized geometry of IP_5_ within the I-site of the mTOR FAT domain features a rotationally shifted mode of binding with a similar number of phosphate/cation contacts as compared to IP_6_. **(C)** Graphical model depicting the proposed role of IP_5_ on mTORC1 signaling.

**Table 1:** I-site binding energies and phosphate contact counts for IP ligands.

| Ligand | Phosphate/cation contacts<sup>a</sup> |  |  |  |  |  |  | Binding Energy<sup>b</sup> (kcal/mol) |
|---|---|---|---|---|---|---|---|---|
|  | C1 | C2 | C3 | C4 | C5 | C6 | total |  |
| IP<sub>4</sub> (I-1,4,5,6-P<sub>4</sub>) | 2 | - | - | 2 | 3 | 2 | 9 | -204.6 |
| IP<sub>5</sub> (I-1,3,4,5,6-P<sub>5</sub>) | 1 | - | 1 | 3 | 2 | 3 | 10 | -232.4 |
| H<sub>2</sub>IP<sub>6</sub> | 1 | 3 | 2 | 0 | 2 | 3 | 11 | -216.3 |
| HIP<sub>6</sub> |  |  |  |  |  |  |  | -239.5 |
| IP<sub>6</sub> |  |  |  |  |  |  |  | -251.4 |

<sup>a</sup> Phosphate groups are labeled by the carbon atom to which they are bonded, with numbering as shown in Figures 6 A and B). Phosphate/cation contacts = number of lysine or arginine residues of the I-site that make close contact the phosphate group. <sup>b</sup>ONIOM (wb97xd/cc-pvtz:wb97xd/6-31g(d)) binding energies were obtained from partially-optimized complexes of the ligand bound to a cluster model of the I-site.

## Discussion

mTORC1 activity plays a central role in regulating cellular metabolism. In pancreatic islets, mTORC1 regulates β-cell mass and function in response to nutrient overload and increased insulin demand (11). While transient mTORC1 activation supports adaptive increases in insulin production, chronic elevation of basal mTORC1 signaling promotes metabolic dysregulation and β-cell dysfunction (12, 16). Despite extensive characterization of the pathways that initiate mTORC1 signaling, the mechanisms that support sustained mTORC1 hyperactivation remain incompletely understood.

Using complementary genetic and pharmacological approaches, we show here that suppression of IPMK and/or ITPK1 reduces cellular IP_5_ levels and selectively impairs basal and insulin-stimulated mTORC1 signaling without disrupting upstream PI3K/Akt or AMPK activation.

Reduced IP_5_ levels did not delay or prevent mTORC1 activation by growth factor or glucose but accelerated termination of the mTORC1 signal following upstream pathway termination. These findings indicate that inositol phosphates are not required for mTORC1 activation per se but instead function to stabilize the active complex and prolong signaling, which is consistent with our previous *in vitro* findings (5). This distinction may be particularly relevant in β-cells exposed to excess nutrients, where elevated basal mTORC1 activity contributes to metabolic acceleration, loss of nutrient responsiveness and hyperinsulinemia (2, 4, 17).

IPMK has been previously implicated in mTORC1 regulation through kinase-independent mechanisms, including scaffolding interactions with Raptor and sequestration of AMPK, as well as through effects on Akt signaling (14, 18, 19). The use of a selective IPMK inhibitor in the present study allowed us to acutely suppress IPMK catalytic activity without altering IPMK protein levels or scaffolding functions. Under these conditions, mTORC1 signaling was impaired despite intact Akt activation, lysosomal recruitment and unchanged AMPK, indicating that IPMK-dependent regulation of mTORC1 cannot be explained by upstream signaling or non-catalytic roles. Importantly, acute inhibition of IPMK kinase activity was sufficient to attenuate mTORC1 signaling within hours, arguing against indirect effects mediated by altered gene expression, cell proliferation, or long-term metabolic adaptations. Furthermore, suppression of ITPK1, which has no known scaffolding functions, phenocopied the effects of IPMK inhibition on IP_5_ levels and mTORC1 signaling. The convergence of these independent perturbations strongly supports a metabolite-dependent mechanism. Together with our previous findings demonstrating that IP_4_, IP_5_, and IP_6_ directly enhance and prolong enzyme activity *in vitro* (5), these data support a model in which higher-order inositol phosphates stabilize active mTORC1 in cells through direct binding to mTOR (Fig. 6C).

Among the inositol phosphate species examined, IP_5_ emerged as the metabolite most closely correlated with mTORC1 signaling. IP_5_ was consistently reduced across all perturbations that impaired mTORC1 activity, whereas IP_6_ levels remained unchanged unless both IPMK and ITPK1 pathways were simultaneously suppressed. Several other observations further support IP_5_ as the primary regulator: (i) the tight association of IPMK with Raptor (18) strongly suggest that IP_5_ is generated near the site of mTORC1 activation; (ii) IP_5_ activates mTORC1 *in vitro* at concentrations comparable to IP_6_ but lower than those required for IP_4_ (5); (iii) computational modeling of IP_5_ binding to the I-site of the mTOR FAT domain shows that optimal contact with the positively charged residues in the pocket can be achieved with 5 phosphates (Fig. 6B and Table I). Together, these findings favor a model in which IP_5_ stabilizes the active conformation of mTORC1, thereby prolonging signaling after activation (Fig. 6C). This could be a potential mechanism by which active mTORC1 can leave the lysosomal surface, where it is initially activated by RHEB, and reach substrates that are not in the vicinity of the lysosome. It could also explain sustained basal (growth-factor-independent) mTORC1 activity in pathological states.

Our results reveal a compensatory relationship between IPMK and ITPK1 in maintaining IP_5_ supply for IP_6_ synthesis. Although inhibition of either enzyme alone substantially reduced IP_5_, IP_6_ levels were preserved, indicating that synthesis of IP_5_ is not the rate limiting step in IP_6_ generation in ß-cells. This metabolic redundancy underscores the selective pressure for maintaining a stable IP_6_ pool, which is known to stabilize the conformation of multiple proteins (20).

Suppression of IPMK or ITPK1 impaired ß-cell metabolic adaptation to elevated glucose, demonstrating the impact of inositol phosphate-dependent mTORC1 regulation on ß-cell function. Future studies will focus on understanding how ITPK1 and IPMK activities impact glucose-dependent insulin secretion and mTORC1-mediated pancreatic ß-cell dysfunction in pathological states such as obesity, diabetes and premature aging.

## Acknowledgements

We are thankful to Dr. John York and Dr. Richard Honkanen for insightful discussions. Computational work was made possible in part by a grant of high-performance computing resources and technical support from the Alabama Supercomputer Authority. This work was supported by the National Institute on Aging, R21 NIA AG071975 to L.E.R.

## Materials and Methods

### Generation of IPMK and ITPK1 knockdown cells

pSuper.retro.puro vectors (OligoEngine) containing the DNA insert for expressing the intended siRNA target sequences for rat IPMK, rat ITPK1 or mutated versions (as control) were designed and constructed according to the manufacturer’s instructions (see Table for siRNA sequences). Vectors were sequenced to confirm insert presence and proper sequence.

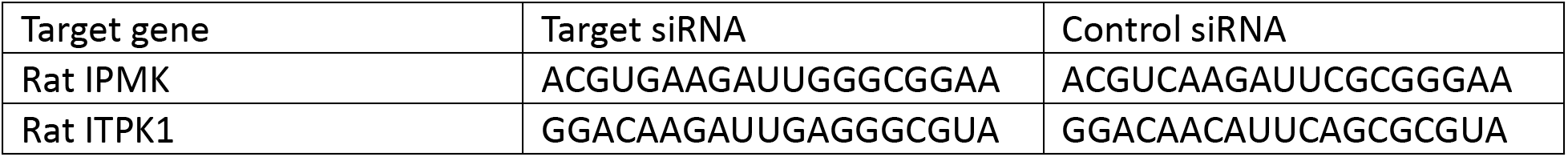

Retrovirus particles for each pSuper construct were packed using HEK293 cells expressing VSVG and gag-pol. Virus particles were collected from the media, filtered and used to infect INS-1 cells in the presence of polybrene. Infected cells were selected using puromycin (0.5 μg/ml). Pools of 100-500 puromycin-resistant colonies per vector were combined for analysis.

### Cell culture

INS-1 832/13 cells (uninfected or infected with pSuper virus as described above) were cultured in RPMI1640 media supplemented with 10% FBS (fetal bovine serum), 10 mM Hepes, 1mM sodium pyruvate and ß-mercaptoethanol in a 37°C incubator with 5% CO_2_. Cells were passage 1:5 every 3 to 5 days.

We previously demonstrated that culturing INS-1 cells in 4.5 mM glucose (which are the normal glucose levels in the circulation) for 2-21 days reduced basal mTORC-1 signaling and improved glucose fold response for mTORC1 stimulation and for insulin secretion (2). Thus, most of the experiments in this study were carried out in INS-1 cells (uninfected or infected with pSuper virus as described above) pre-cultured in low glucose containing media (glucose-free RPMI1640 media supplemented with 4.5 mM glucose, 10 mM Hepes, 1mM sodium pyruvate, 10% FBS and ß-mercaptoethanol) for 3 days prior to the start of each experiment, unless indicated.

### Human Islets

Non-diabetic human Islets were obtained through the services of the Integrated Islet Distribution Program (IIDP). Islets were cultured in non-treated tissue culture plates with RPMI media containing 5.5 mM glucose and supplemented with 10% FBS (fetal bovine serum), 10 mM Hepes and 1mM sodium pyruvate in a 37°C incubator with 5% CO_2_.

### qPCR analysis

mRNA from INS-1 cells were collected using QIAwave RNA plus mini kit (Qiagen). cDNAs were prepared using Verso cDNA kit and 0.5 μg mRNA. qPCR reactions were performed with SYBRGreen reagents, specific primer sets (IDT) according to Table II, and the cDNA reaction (1:800 dilution). Quantitative real time PCR reactions were carried using CFX Connect real time PCR system (Bio Rad).

### Measurements of cellular IP levels

HEK293: Cells were plated in a 6-well poli-L-lysine coated plate in DMEM-10% FBS. The next day, cells were metabolically labeled with [^3^H]-inositol (Revvity, Inc) at a concentration of 25 μCi/ml in DMEM-10% FBS. 48hrs later the labeling media was removed and cells were treated with DMSO or 10 μM UNC9750 in DMEM-1% FBS for 5 hrs. Cells were lysed and inositol phosphates were collected and analyzed as described below.

INS-1: Cells pre-cultured in 4.5 mM glucose for 3 days (see above) were plated in a 6-well plate at a density of 2-2.5 ×10^6^ cells per well. The next day, cells were metabolically labeled with [^3^H]-inositol (Revvity, Inc) at a concentration of 25 μCi/ml in MEM (5.5 mM glucose) media supplemented with 10% FBS and Hepes. For UNC9750 treatments, the compound was added for a final of 10 μM concentration, either 1 hr before addition of the [^3^H]-inositol or 19 hrs after, as indicated for each experiment. When indicated, glucose was added to the MEM (which contains 5.5 mM glucose) for a final concentration of 11 mM. After 24 hrs in [^3^H]-inositol, cells were lysed and inositol phosphates were collected and analyzed as described below.

Inositol phosphate analysis: After labeling and treatments, cells were washed with PBS and lysed with 200 μl HCl (0.5 M), 250 μl methanol and 125 μl KCL (2M). The lysates were scraped from the plates, sonicated (3 one-minute cycles with a cup-horn sonicator at 4°C) and purified from proteins and lipids through 2 consecutive chloroform extractions. The water-soluble phase was dried in a speedvac concentrator, resuspended in 10 mM ammonium phosphate. Labeled inositol phosphate species were separated by anionic exchange HPLC (0.03-1.3 M ammonium phosphate in 60 minutes) and analyzed by an on-line flow scintillation analyzer. The peaks corresponding to cellular IP_4_, IP_5_, IP_6_ and IP_7_ were identified using [^32^P]-labeled IP_4_, IP_5_, IP_6_ and IP_7_ generated from *in vitro* reactions with purified IPMK (for IP_4_ and IP_5_), IPK1 (for IP_6_) or IP6K (for IP_7_) enzymes. After quantification of the cpm in each peak, the data were normalized against the cpm present in the [^3^H]-inositol peak.

### Western-blot and Analysis of mTORC1 signaling in INS1, HEK293 and human islets

HEK293: Cells were platted into poli-L-lysine coated 24-well tissue culture dishes in DMEM media supplemented with 10% FBS. After 24 hrs, the media was replaced with DMEM containing 1% FBS. The next day, cells were treated with DMSO or UNC9750 at 10 μM for 2-3 hrs and stimulated with IGF-1 (10 nM) or PBS control for 30 minutes before protein lysates were collected, as described below.

INS-1: After pre-culture in low glucose (4.5-5 mM) for 3 days, cells were platted into 24-well tissue culture dish coated with poli-L-lysine at a density of 350,000 to 500,000 cells per well. 48 hrs after platting, cells were placed in serum-free media (RPMI1640 containing 4.5 mM glucose, 10 mM Hepes, 1mM sodium pyruvate) for 2 hrs. 10 nM insulin was added to the rest media in some experiments, as indicated. For UNC9750 experiments, the compound was added at a concentrations of 5-10 μM, during the rest period, unless otherwise indicated. After the 2 hrs rest in serum-free media, cells were stimulated with insulin or glucose according to each experimental description. For mTORC1 termination experiments, cells were treated with insulin (100 nM) for 45 minutes, after which wortmannin (100 nM) or rapamycin (20 nM) was added.

Human islets: After three days in culture, islets were dispersed using a mixture of Versene (Gibco) and trypsin (4:1). Cells were plated on a poli-L-lysine coated low area 96-well plate, at about 50,000 cells/well in RPMI media containing 5.5 mM glucose and supplemented with 10% FBS (fetal bovine serum), 10 mM Hepes and 1mM sodium pyruvate. 2-3 days later, the media was replaced with RPMI containing 4.5 mM glucose, 10 mM Hepes and 1mM sodium pyruvate. Cells were treated with DMSO or UNC9750 for 2 hrs and stimulated with control PBS, 10 nM IGF-1 or 12 mM glucose for 1 hr before protein lysates were collected, as described below.

Protein lysates were collected using denaturing lysis buffer [50 mM Tris-HCl (pH 7.5), 100 mM NaCl, 1 mM EDTA, 1% Triton-X100, 1.5 % SDS, 100 mM DTT and 10% glycerol] containing protease inhibitor cocktail (Sigma) and phosphatase inhibitors (1 mM sodium orthovanadate, 2 mg/ml sodium fluoride and 2 mg/ml ß-glycerophosphate). Lysates were shaken for 30 minutes at 1400 rpm and 65°C to mechanically disrupt the DNA. Proteins were resolved by SDS-PAGE (polyacrylamide gel electrophoresis) using pre-cast 4-15% gradient polyacrylamide mini gels (Bio-Rad) and transferred to nitrocellulose. For p70S6K mobility shift assays, lysates were run in a 10 cm SDS-PAGE using lab-casted 10% polyacrylamide gel. After blocking the membranes with 5% non-fat milk and incubating with primary antibody overnight, bands were visualized using IR-680 or IR-800 conjugated secondary antibodies and fluorescence was detected using Typhoon infrared scanner (Cytiva) and quantified using ImageStudio software (LiCor). Antibodies against pS6 (S235/236), total S6, pAMPK (T172), pAkt (S473), ITPK1, actin, tubulin, p70S6K were from Cell Signaling Technologies, antibody against IPMK was from Novus biological; rabbit IR680 was from Li-Cor and mouse IR800 were from Cytiva.

P values were calculated using Student’s t-test (two tail distribution, two samples equal variance). For pancreatic islet analysis we also used two-way ANOVA to compare DMSO-and UNC970-treated groups.

### mTORC1 complex analysis

Cells were cultured in 100 mm tissue culture dish for 48 hrs as described above, treated with DMSO or 10 μM UNC9750 for 2-3 hrs before cell lysis in CHAPS lysis buffer (40 mM Hepes, 40 mM NaCl, 1 mM EDTA, 0.3% CHAPS) containing protease inhibitor cocktail (Sigma) and phosphatase inhibitors (1 mM sodium orthovanadate, 2 mg/ml sodium fluoride and 2 mg/ml ß-glycerophosphate). Lysates were cleared by centrifugation at 14,000 rpm for 10 minutes, incubated with primary antibody at 4°C overnight and with protein A/G agarose (blocked with 5% BSA) for 2 hrs. Beads were washed 4 times with lysis buffer, mixed with SDS-containing loading buffer and incubated at 95°C for 5 minutes. Proteins present in the immunocomplex were resolved by SDS-PAGE (polyacrylamide gel electrophoresis) using pre-cast 4-15% gradient polyacrylamide mini gels (Bio-Rad), transferred to nitrocellulose and processed as described for Western-blots above. For mTOR immunoprecipitation, anti-mTOR (CST#2983) was used and for Raptor (CST#2280). P values were calculated using Student’s t-test (two tail distribution, two samples equal variance).

### mTOR subcellular localization

HEK293 cells were plated into poli-L-lysine coated glass bottom 12-well plates in DMEM-10%FBS. 24 hrs after plating, cells were cultured in DMEM-1% FBS overnight, treated with DMSO or 10 μM UNC9750 for 2-3 hrs and with PBS or 10 nM IGF-1 for 30 minutes. Cells were fixed in PBS-4% para-formaldehyde for 5 minutes, permeabilized in PBS-0.2% Triton X-100 for 15 minutes, blocked with PBS-0.25% BSA or PBS-10% donkey serum for 1 hr and incubated with primary antibodies (diluted in blocking buffer) at 4°C overnight. After 3 washes in PBS, cells were incubated with secondary antibody for 1 hr at room temperature. For mTOR staining, rabbit anti-mTOR (CST#2983) was used and for Lamp1 staining, mouse anti-Lamp1 (CST#15665) was used. Secondary antibodies were Donkey anti-Rabbit conjugated with Cy5 and Donkey anti-mouse conjugated with Cy2 (Jackson Laboratories).

Images were acquired with a Nikon AXR-NSPARC laser scanning confocal microscope, using a Plan Apo λD 20x Plan Apo (NA=0.8) objective, and operating in galvanometer-scanning super-resolution mode (pixel resolution =.19 um). Image analysis was performed using an image analysis pipeline implemented in Python to measure the distribution and colocalization of the lysosomal and mTOR signals on a per-cell basis. To segment cells, binary masks were first generated by Guassian smoothing and Otsu thresholding of the DAPI and mTOR channels to generate nuclear and total-cell masks, respectively. Segmented nuclei were then labeled and expanded until they touched neighboring labels without overlap. Individual cells were then defined as the intersection of each expanded nuclear label with the total-cell mask, and a cytoplasmic mask for each cell was generated by removing the original nuclear mask. Total lysosomal and mTOR masks were generated using a fixed threshold value (30 and 75 for Lamp1 and mTOR, respectively), and cell-specific masks were generated by intersecting each cell’s cytoplasmic mask with the lysosome and mTOR total masks. All measurements were performed within the individual cell masks. For each cell, lysosomal and mTOR fluorescence intensities were measured, including mTOR intensity within the lysosomal mask. Lysosome–mTOR association was quantified using the Manders colocalization coefficient with the lysosomal mask, Manders overlap coefficient, Pearson correlation coefficient, and the intersection coefficient between the cell-specific lysosomal and mTOR masks. P values were calculated using one-way ANOVA with Turkey’s multiple comparison test.

### Respirometry

The XF^e^24 Extracellular Flux Analyzer (Seahorse Bioscience) was used to measure oxygen consumption rate (OCR). INS-1 cells were seeded in XF24 cell culture plates pre-coated with poly-L-lysine at 60,000 to 80,000 cells per well in complete media containing either 4.5 or 11 mM glucose, according to each experiment. For IPMK inhibition, UNC9750 or DMSO were added at the time of platting, as indicated. 48 hrs after plating, cells were washed in Seahorse base media (Li-Cor) supplemented with 3 mM glucose, 2 mM glutamine and 1 mM sodium pyruvate and incubated in a 37°C incubator without CO_2_ for 2 hrs prior to analysis (resting period). Basal oxygen consumption at 3 mM glucose was measured prior to sequential injection of glucose (8 mM final), followed by oligomycin (5 µM), FCCP (1 µM) and antimycin A (5 µM). At the end of the analysis, cells were fixed with formaldehyde, stained with dapi and the total cell number in each well was determined using Celigo Imaging Cytometer, for the purpose of normalization. Basal respiration was calculated by obtaining the difference between baseline OCR (last point taken before injection of additional glucose) and non-mitochondrial respiration (minimum OCR following antamycin A treatment). Proton leak was calculated by obtaining the difference between the oligomycin-resistant OCR (minimum OCR following oligomycin treatment) and non-mitochondrial respiration. ATP-coupled respiration was calculated by obtaining the difference between basal respiration and proton leak. Maximal respiratory capacity was calculated by obtaining the difference between the uncoupled OCR (the maximum OCR following FCCP treatment) and non-mitochondrial respiration. Percentage leak (mitochondrial inefficiency) was calculated relative to basal respiration after glucose addition. P values were calculated using Student’s t-test (two tail distribution, two samples equal variance).

### Computational Modeling

Quantum calculations using Gaussian16 (21) were carried out to rank orientations of IP_5_ and IP_4_ within the I-site of mTOR’s FAT domain and to obtain I-site binding energies. First, a 13-residue cluster model of the I-site was built (charge = +8) from the cryo-EM structure of mTORC2:IP_6_ (pdb entry 6ZWO; 3.0 Å res. (6)). Initial IP_5_/IP_4_ poses were inferred from the pose of IP_6_ in the solved complex, using six possible rotationally-shifted superpositions of the inositol ring. The IPs were modeled as chairs, with the C2 hydroxyl in an axial position, because at present, all reported structures of IP_5_ or IP_4_ in complex with proteins (e.g., pdb entry 4O4E; 1.90 Å res. (22); pdb entry 4A69; 2.06 Å res. (23)) show the inositol ring in this conformation.

Next, the complexes were optimized using the ONIOM hybrid computational model (reviewed in (24)) with quantum methods assigned to both levels of a two-region system. Implicit water was included using the solvent reaction field method (SCRF). During optimizations, the ligand and contacting sidechains of the pocket were free to move, while the rest of the system was frozen. IP_5_ and IP_4_ were modeled as fully deprotonated species (charge = −10 and −8, respectively). In the first screen of orientations, the wb97xd density functional and 6-31g(d) basis set were used for the high-level, while the PM7 semi-empirical method was used for the low-level; the second screen instead used the cc-pvtz basis set for the high-level region.

For comparison, optimizations of the IP_6_ complex, with IP_6_ in different protonation states (H_2_IP_6_: H^+^ on C4 and C1 phosphates; HIP_6_: H^+^ on C4 phosphate; and IP_6_; charge = −10, −11, and −12, respectively) were carried out. After optimization of the H_2_IP_6_ complex, we observed that the initial orientations of K1788 and K1662 taken from the cryo-EM structure could be adjusted to improve phosphate contacts. Such adjustments were preset for the HIP_6_ and IP_6_ cases and were also applied to follow-up re-optimizations of the H_2_IP_6_ complex and the best IP_5_ and IP_4_ complexes. Binding energies, computed as E(complex) – E(free ligand) – E(empty I-site), were assigned after final ONIOM(wb97xd/cc-pvtz:wb97xd/6-31g(d)) optimizations met default thresholds for RMS and maximum forces.

Figures 6 A and B were generated using Discovery Studio Visualizer v24.1.0.23298 (Dassault Systemes Biovia Corp 2023). Optimized structures, including those of free ligands (solvated), are provided in the Supplemental Materials, along with Gaussian input files in which the atoms of the high/low regions and frozen/free regions are indicated.

## Notes

### Competing Interest Statement

X.W. is founder, equity holder, and serves as a scientific advisor for InoKare Therapeutics, Inc..

### Summary of Updates

New data showing the effect of IPMK inhibition on mTORC1 signaling in HEK293 and human pancreatic islet cells. New data showing the lack of effect of IPMK inhibition on mTORC1 complex formation or mTOR colocalization with Lamp1. New data showing the effect of IPMK or ITPK1 suppression on cellular metabolism in response to chronic excess glucose. Two new authors were included.

