## Supplemental Figures for "Suppression of ITPK1 and IPMK activities impairs mTORC1 signaling due to accelerated signal termination and prevents pancreatic ß-cell metabolic adaptation to excess glucose"

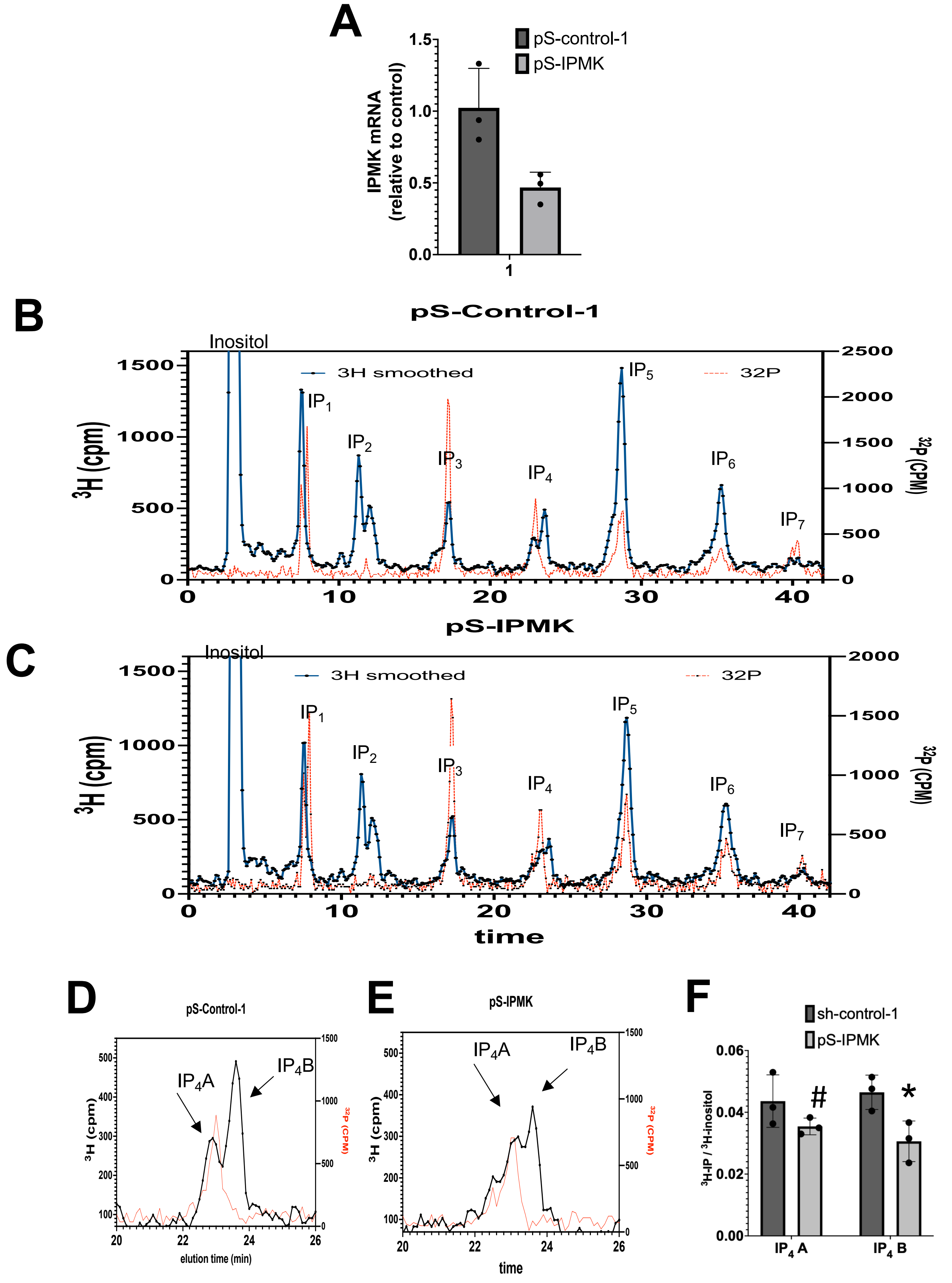


Supplemental Figure 1

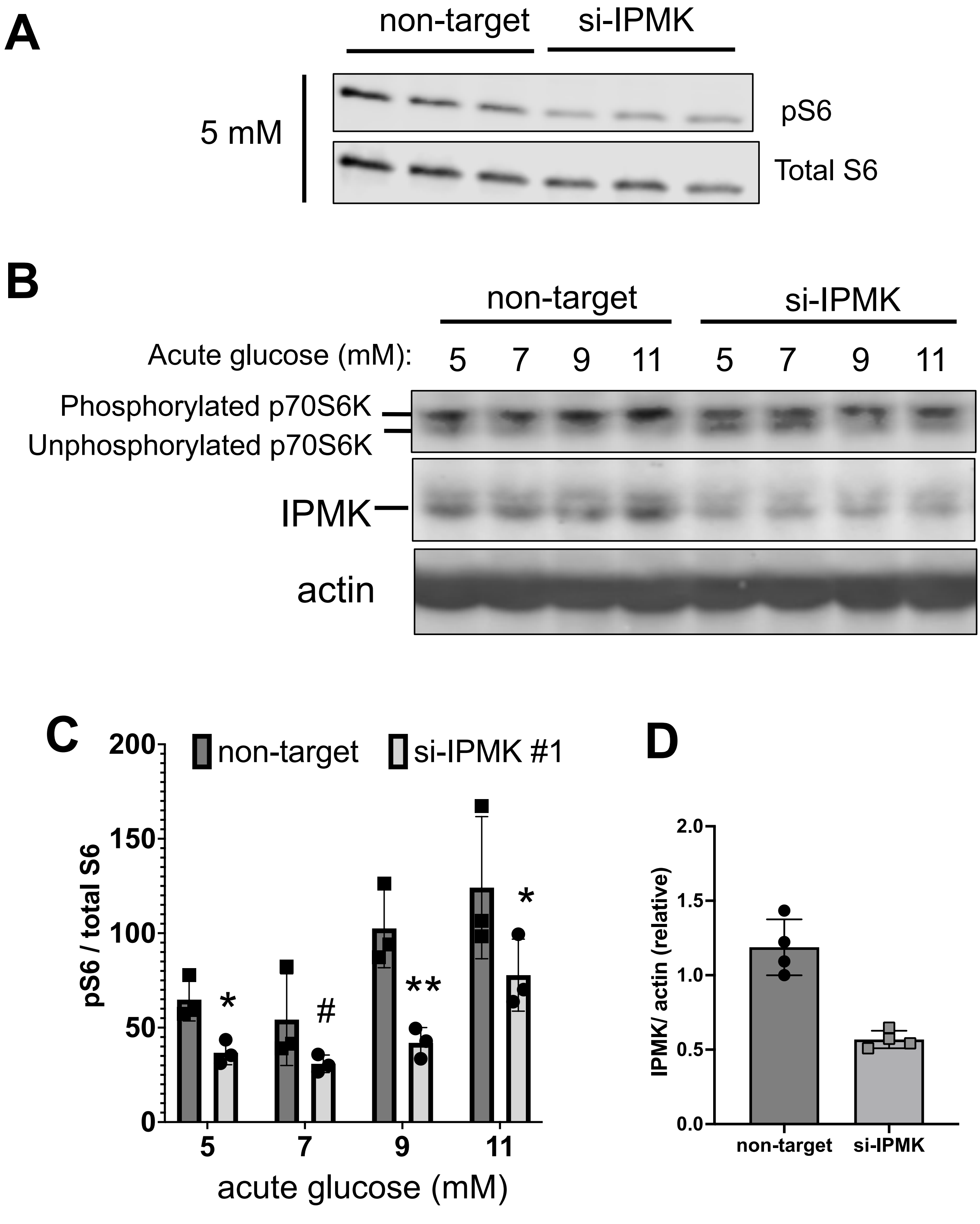


Supplemental Figure 2

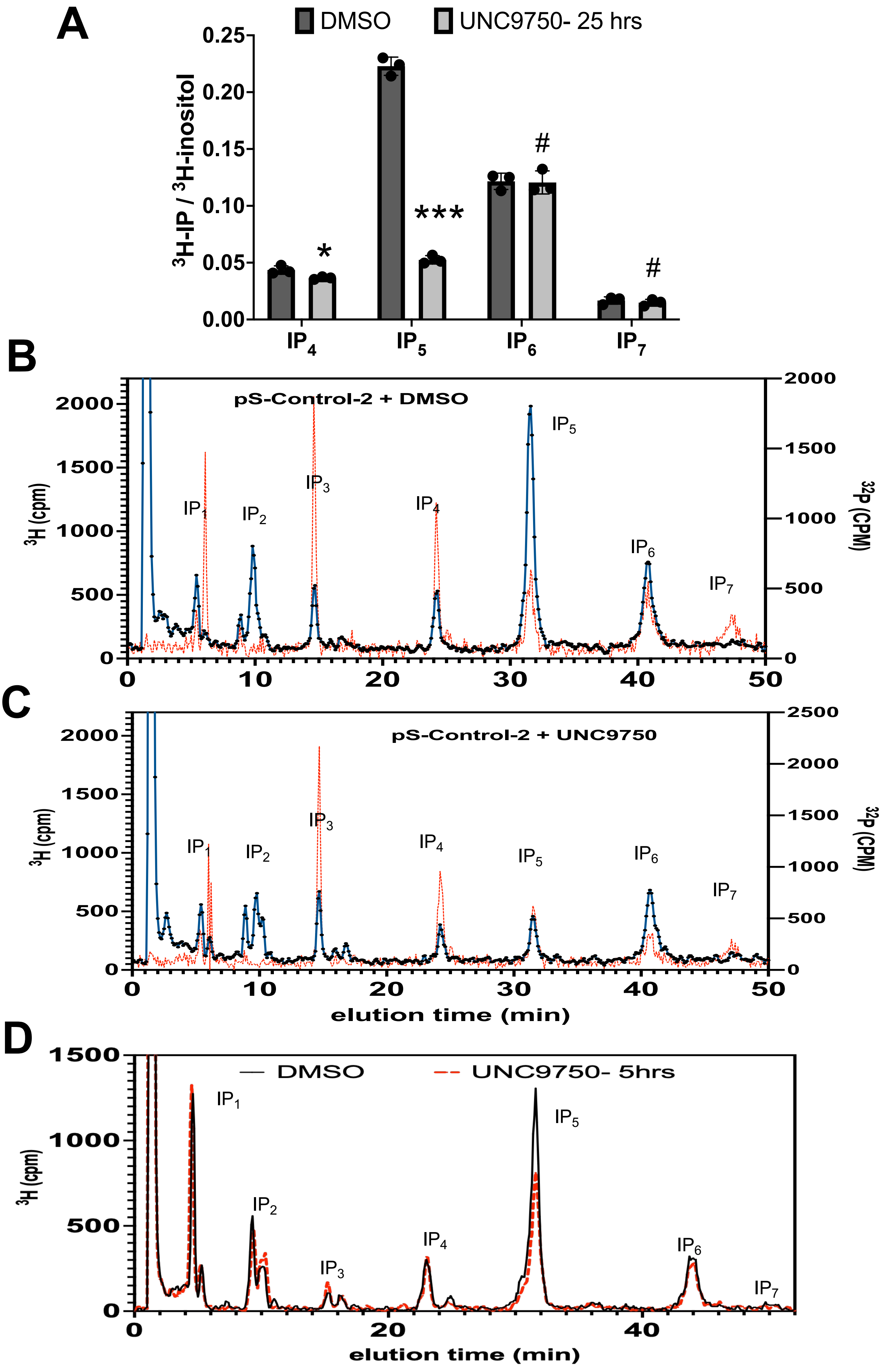


Supplemental Figure 3

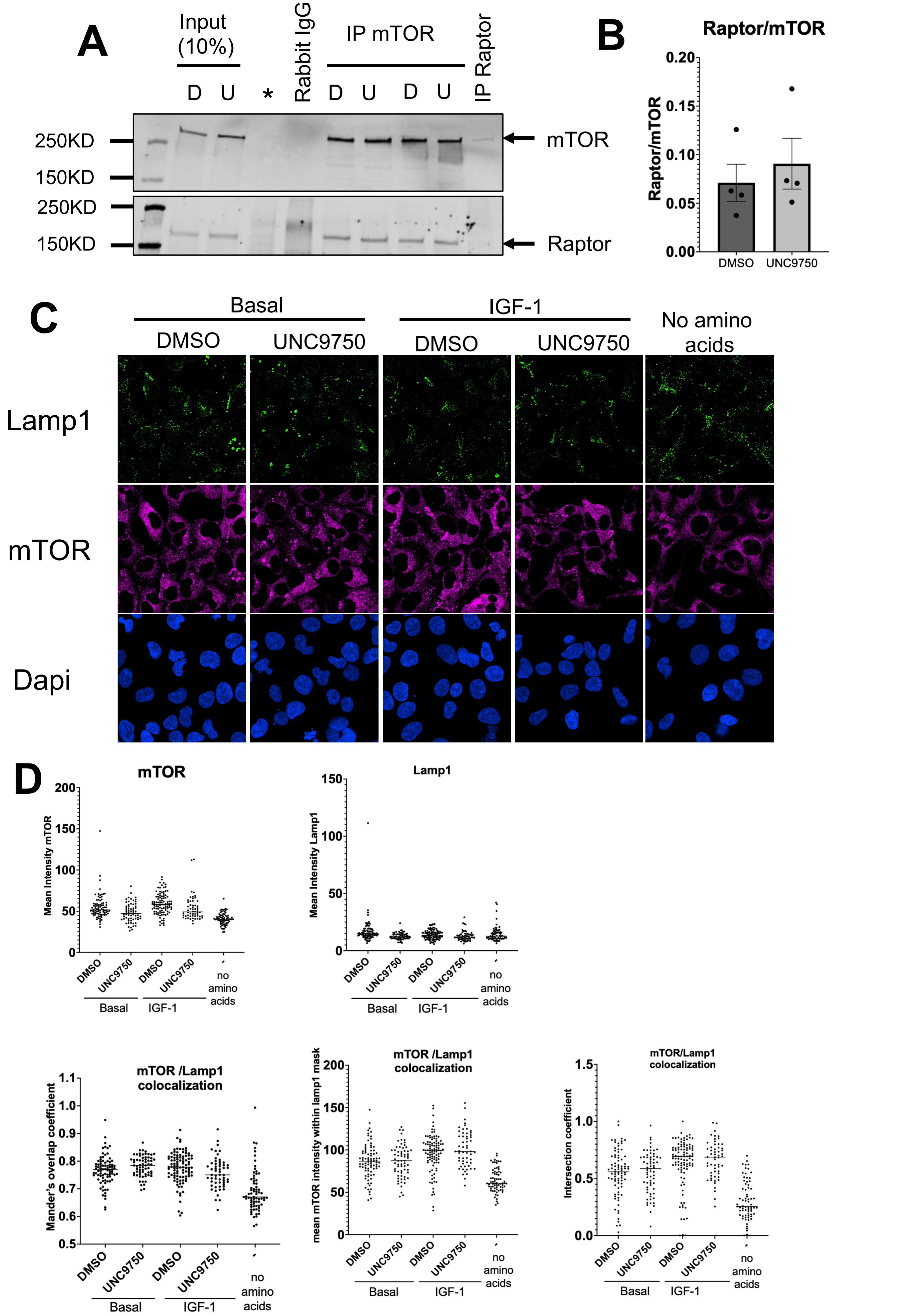


Supplemental Figure 4

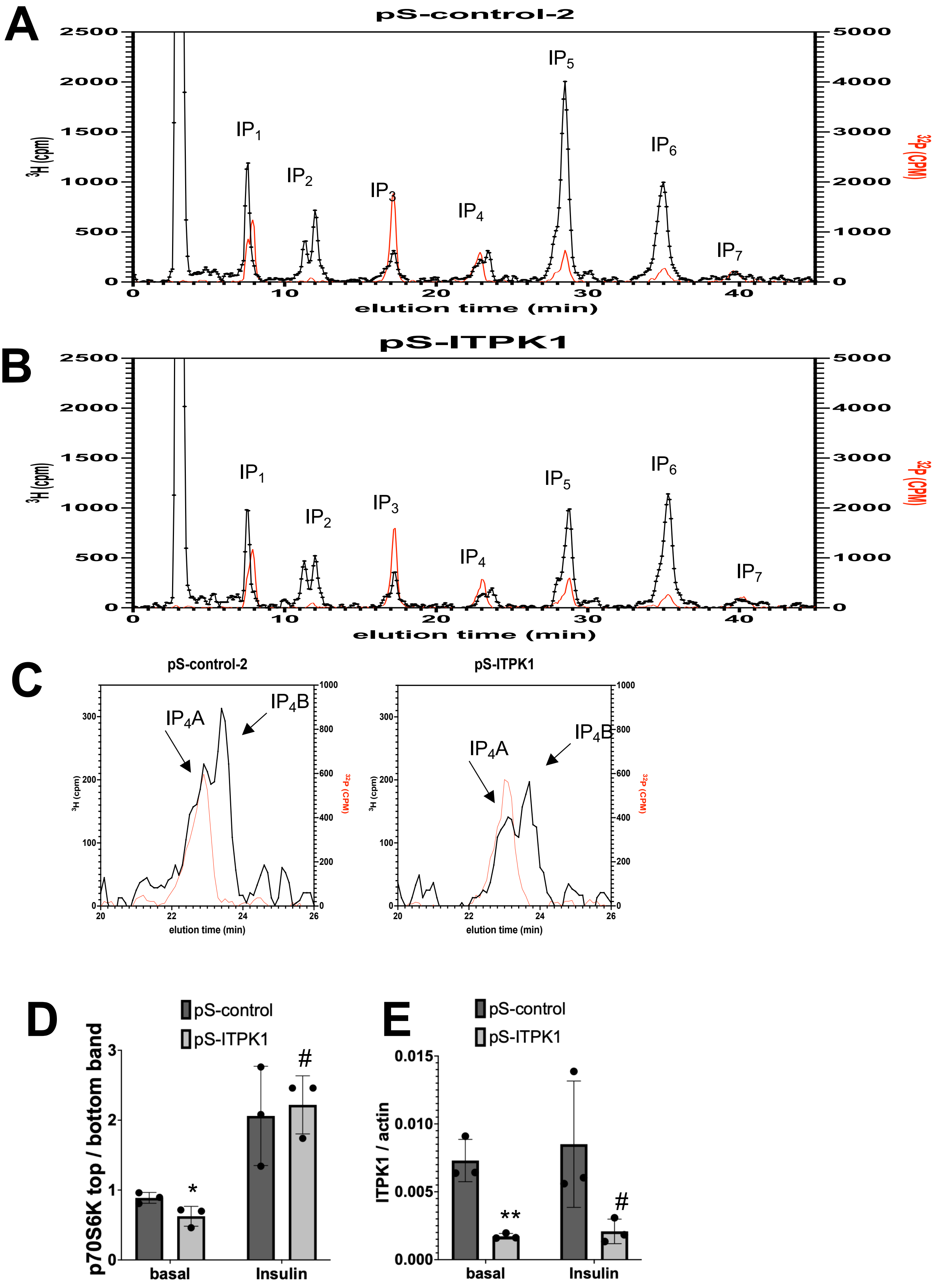


Supplemental Figure 5

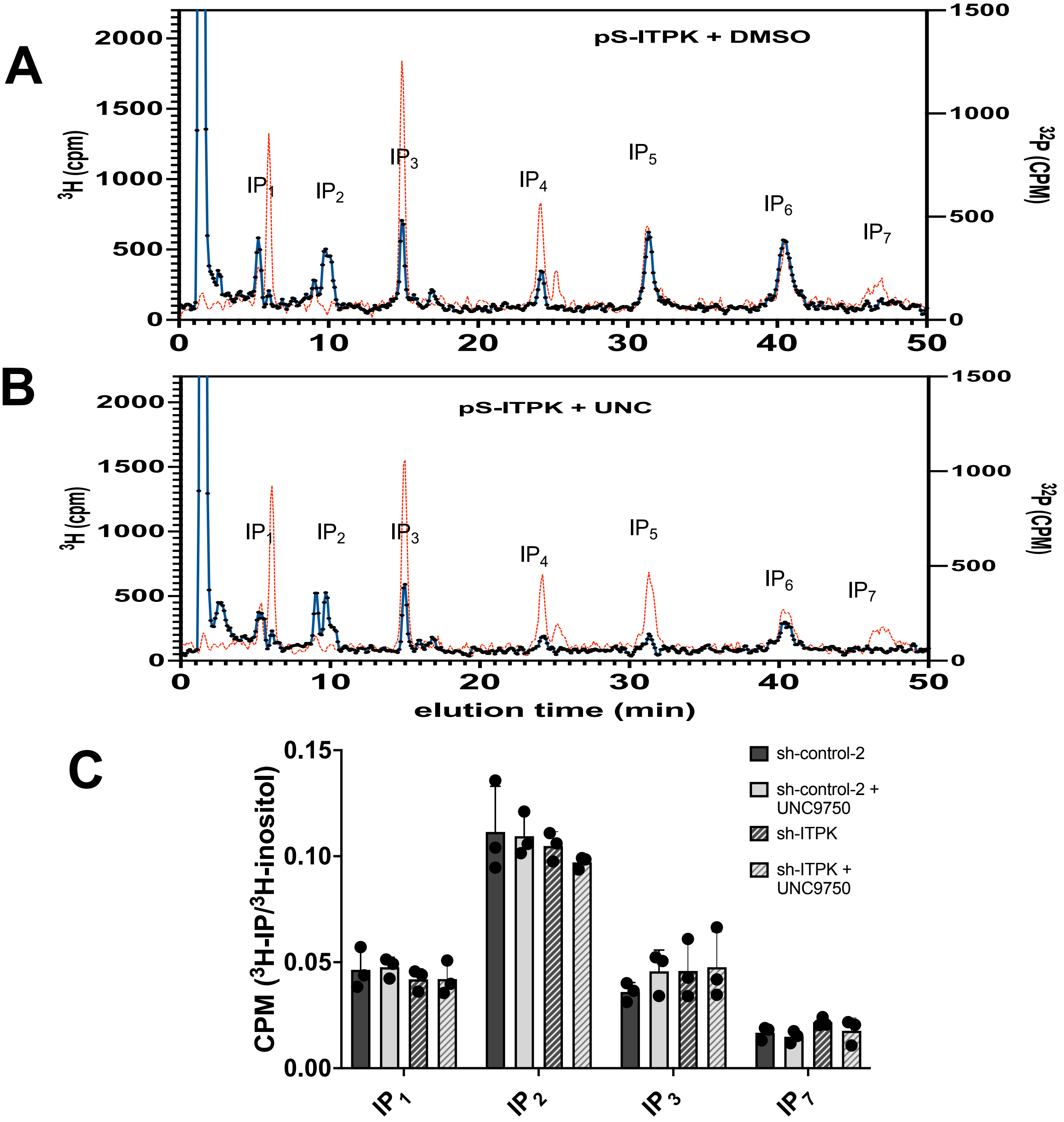


Supplemental Figure 6

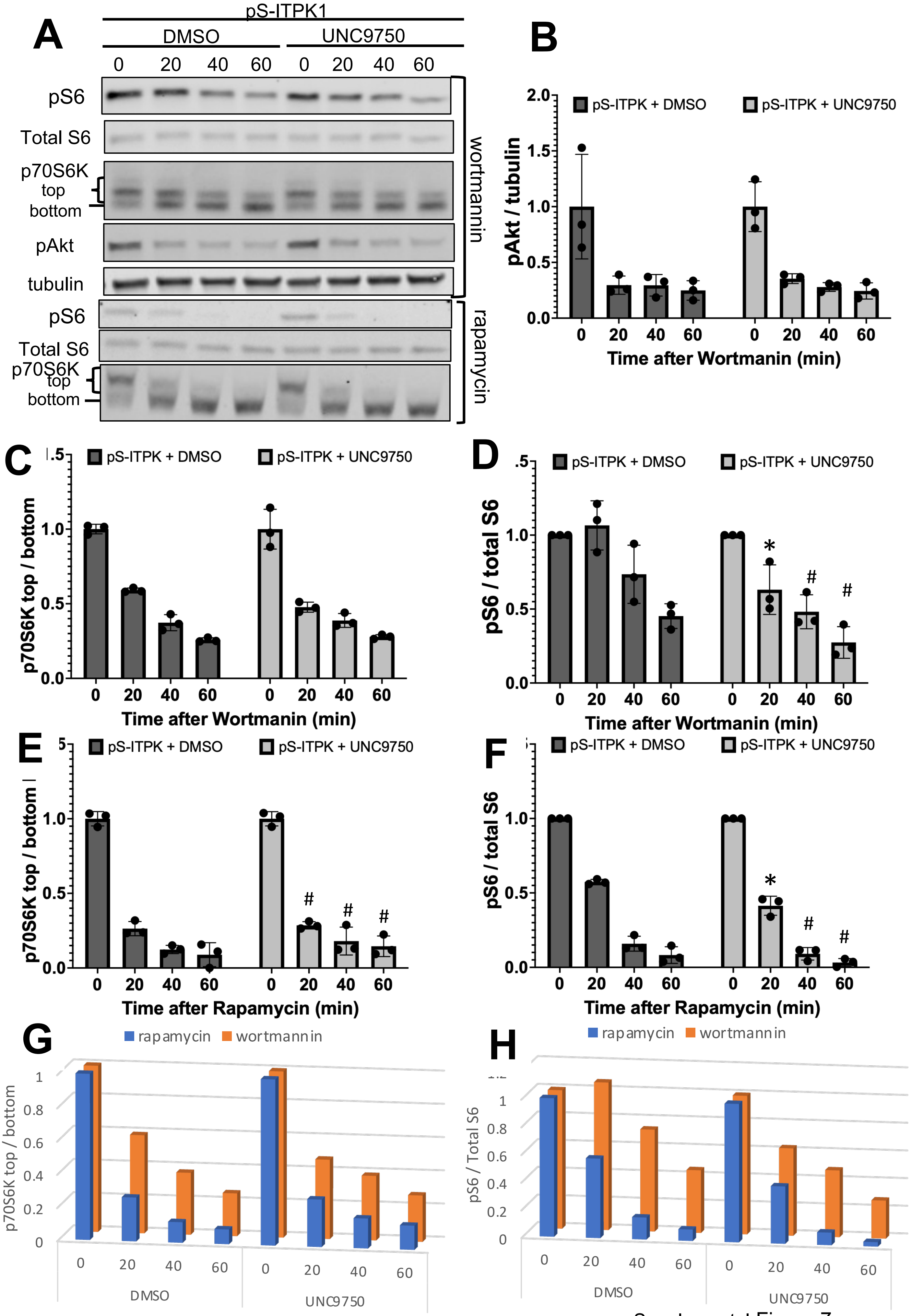


Supplemental Figure 7

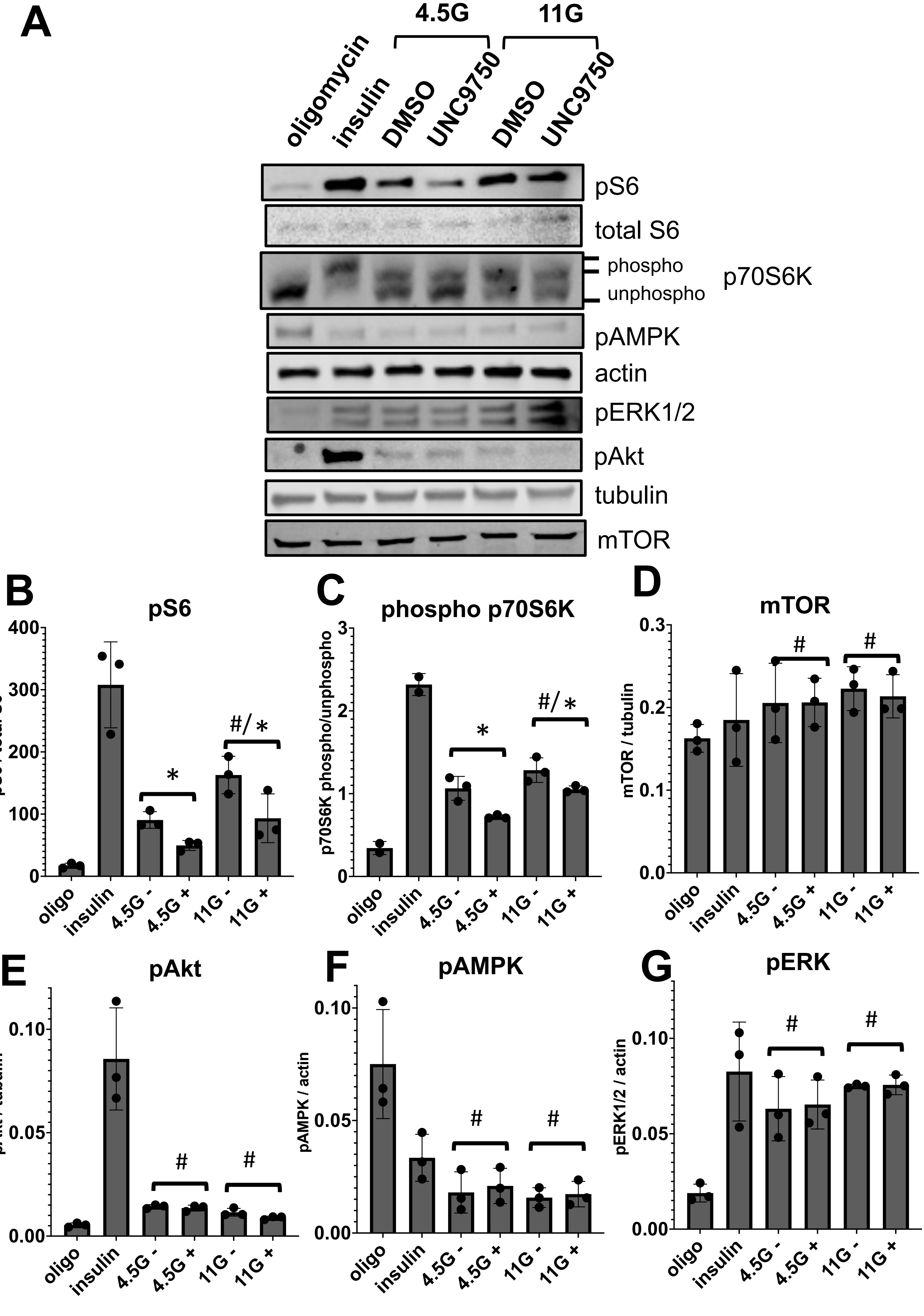


Supplemental Figure 8
