## Supplemental Figure Legends for "Suppression of ITPK1 and IPMK activities impairs mTORC1 signaling due to accelerated signal termination and prevents pancreatic ß-cell metabolic adaptation to excess glucose"

**Figure S1: IPMK knockdown reduced $IP_4$ and $IP_5$, without affecting $IP_3$, $IP_6$ or $IP_7$.**

**(A)** IPMK mRNA in INS-1 cells infected with pS-control-1 or pS-IPMK, as measured by qPCR. Data shown are the mean and standard deviation of 3 biologically independent samples.

**(B-C)** Representative HPLC elution profile of cellular [$^3$H]-inositol labeled peaks (blue lines) from INS-1 with pS-control-1 (B), pS-IPMK (C). The dotted red lines show the [$^{32}$P] channel which contained internal standards for $IP_4$, $IP_5$, $IP_6$ and $IP_7$ (red lines). Note that the [ $^{32}$P]-peaks comigrating with $IP_1$, $IP_2$ and $IP_3$ are breakdown products from the higher order IPs labeled with $^{32}$P.

**(D-E)** Same as in (B-C) except that only the region for $IP_4$ peaks is shown. The dotted red lines show the [$^{32}$P] channel which contained internal standards for $IP_4$ generated using purified IPMK.

**(F)** Quantification of the [$^3$H]-$IP_4$ subpeaks A and B (as indicated in D and E) from INS-1 cells infected with pS-control-1 or pS-IPMK. Data shown are the mean and standard deviation of 3 biologically independent samples normalized for [$^3$H]-inositol. (*) P=0.03; (#) P>0.05.

**Figure S2**: **Transient IPMK knockdown impaired mTORC1 signaling.**

**(A)** Representative western blots showing the effects of transient IPMK knockdown on mTORC1 signaling as measured by S6 phosphorylation (pS6) in INS-1 cells transiently transfected with non-target si-RNA or si-RNA targeting IPMK (si-IPMK) and kept in media with low glucose (5 mM).

**(B)** Representative western blots showing the effects of transient IPMK knockdown on mTORC1 signaling as measured by p70S6K mobility shift (top bands are the phosphorylated form and bottom is the unphosphorylated form) and on IPMK protein levels. Cells were treated as in (A) and stimulated for 30 minutes with various glucose concentrations, as indicated.

**(C)** Mean and standard deviation of pS6 levels from 3 biologically independent samples treated as described in B and normalized by total S6. (*) P=0.013-0.02; (**) P=0.009; (#) P>0.05.

**(D)** Mean and standard deviation of IPMK protein from 3 biologically independent samples treated as described in B and normalized by actin.

**Figure S3**: **IPMK inhibition reduced $IP_4$ and $IP_5$, without affecting $IP_3$, $IP_6$ or $IP_7$.**

(**A**) Cellular inositol phosphate levels in INS-1 cells treated with vehicle control (DMSO) or UNC9750 for 25 hrs and normalized to [$^3$H]-inositol. IPMK inhibition reduced $IP_5$ and $IP_4$, with no significant change in $IP_6$ or $IP_7$ **.**

**(B-C)** Representative HPLC elution profile of cellular [$^3$H]-inositol labeled peaks (blue lines) from INS-1 treated with DMSO (A) UNC9750 (B) for 25 hrs. The dotted red lines show the [$^{32}$P] channel which contained internal standards for $IP_4$, $IP_5$, $IP_6$ and $IP_7$ (red dotted lines). Note that the [ $^{32}$P]-peaks comigrating with $IP_1$, $IP_2$ and $IP_3$ are breakdown products from the higher order IPs labeled with $^{32}$P.

**(D)** Representative HPLC elution profile of cellular [$^3$H]-inositol labeled peaks from INS-1 cells treated with DMSO (black line) or UNC9750 (red dotted line) for 5 hrs.

**Figure S4: IPMK inhibition does not affect mTOR/Raptor complex or mTOR co-localization with lysosome.**

**(**A) Representative western blots showing mTOR or Raptor bands present in anti-mTOR, anti-Raptor or control IgG immunoprecipitates (IP) from INS-1 cells pre-treated with control DMSO (D) or UNC9750 (U) for 3 hrs. Also shown are mTOR and Raptor present in the input lysates.

**(B)** Quantification of the Raptor band present in anti-mTOR IP from INS-1 cells pre-treated with control DMSO (D) or UNC9750 (U) for 3 hrs, normalized by mTOR. Data are the mean and standard deviation from 4 separate IPs.

**(C)** Immunofluorescence images of HEK293 cells treated with DMSO or UNC9750 and stimulated or not with IGF-1. mTOR staining is shown in magenta, Lamp1 in green and Dapi in blue. Also shown are control cells starved for amino acids for 30 minutes. Images shown are from the same fields shown in Fig 2F.

**(D)** Quantification of mTOR and Lamp1 from the same experiments shown in Fig 2F and S4 C, and mTOR/Lamp1 co-localization as measured by Mander's overlap coefficient, mean mTOR intensity within Lamp1 mask, and intersection coefficient.

**Figure S5: ITPK1 knockdown cells reduced $IP_4$ and $IP_5$, without affecting $IP_3$, $IP_6$ or $IP_7$.**

**(A-B)** Representative HPLC elution profile of cellular [$^3$H]-inositol labeled peaks (black lines) from INS-1 with pS-control-2 (A), pS-ITPK1 (B). The dotted red lines show the [$^{32}$P] channel which contained internal standards for $IP_4$, $IP_5$, $IP_6$ and $IP_7$ (red lines). Note that the [ $^{32}$P]-peaks comigrating with $IP_1$, $IP_2$ and $IP_3$ are breakdown products from the higher order IPs labeled with $^{32}$P.

**(C)** Same as in (A-B) except that only the region for $IP_4$ peaks is shown. The dotted red lines show the [$^{32}$P] channel which contained internal standards for $IP_4$ generated using purified IPMK.

**(D)** Quantification of p70S6K phosphorylation (top bands/lower band) from the same experiment as in Fig. 3B.

**(E)** Quantification of ITPK1 band normalized against tubulin, from the same experiment as in Fig 3B.

Data shown are the mean and standard deviation of 3 biologically independent samples (*) P=0.05; (**) P=0.0035; (#) P>0.05.

**Figure S6: IPMK inhibition in ITPK1 knockdown cells reduced $IP_4$ , $IP_5$ and $IP_6$, without affecting $IP_3$ or $IP_7$.**

**(A-B)** Representative HPLC elution profile of cellular [$^3$H]-inositol labeled peaks (solid blue lines) from INS-1 with pS-ITPK1 treated with DMSO (A) or UNC9750 (B) for 24 hrs. The dotted red lines show the [$^{32}$P] channel which contained internal standards for $IP_4$, $IP_5$, $IP_6$ and $IP_7$ (red lines). Note that the [ $^{32}$P]-peaks comigrating with $IP_1$, $IP_2$ and $IP_3$ are breakdown products from the higher order IPs labeled with $^{32}$P.

**(C)** Quantification of the [$^3$H]-$IP_1$, -$IP_2$, -$IP_3$ and -$IP_7$ peaks from INS-1 cells infected with pS-control or pS-ITPK and treated with DMSO or UNC9750, as indicated. Data shown are the mean and standard deviation of 3 biologically independent samples normalized for [$^3$H]-inositol. Note that this quantification is from the same experiment as shown in Fig. 3E. No statistical significance was reached between these measurements.

**Figure S7**: **IPMK inhibition in ITPK1 knockdown cells accelerates the decay of mTORC1 signaling.**

**(A)** Representative western blots showing the decline in mTORC1 signaling after signal termination with wortmannin or rapamycin in ITPK1 knockdown (pS-ITPK1) INS-1 cells treated with DMSO or UNC9750, assessed by p70S6K mobility shift or S6 phosphorylation (pS6), and upstream insulin signaling assessed by Akt phosphorylation.
**(B)** Quantification of Akt phosphorylation from three biologically independent samples, normalized to tubulin. Shown are the rate of signal decay after **wortmannin** treatment normalized by time 0 for each treatment group.

**(C)** Quantification of p70S6K phosphorylation from three biologically independent samples, as a ratio between upper phosphorylated bands over bottom, unphosphorylated band. Shown are the rate of signal decay after **wortmannin** treatment normalized by time 0 for each treatment group.
**(D)** Quantification of S6 phosphorylation from three biologically independent samples, normalized to total S6. Shown are the rate of signal decay after **wortmannin** treatment normalized by time 0 for each treatment group.
**(E)** Quantification of p70S6K phosphorylation from three biologically independent samples, as a ratio between upper phosphorylated bands over bottom, unphosphorylated band. Shown are the rate of signal decay after **rapamycin** treatment normalized by time 0 for each treatment group.
**(F)** Quantification of S6 phosphorylation from three biologically independent samples, normalized to total S6. Shown are the rate of signal decay after **rapamycin** treatment normalized by time 0 for each treatment group.
**(G)** 3-D graph showing the juxtaposition of graphs C (wortmannin treatment in orange) and E (rapamycin treatment in blue).
**(H)** 3-D graph showing the juxtaposition of graphs D (wortmannin treatment in orange) and F (rapamycin treatment in blue).
Data are mean ± SD. Statistical significance is indicated in panels. (*)P=0.01-0.05; (**)P=0.005; (#) P>0.05.

**Figure S8**: **IPMK inhibition impaired mTORC1 signaling without affecting the phosphorylation levels of Akt, AMPK or ERK and did not prevent glucose-induced mTORC1 stimulation**.

**(A)** Representative western blots showing the effects of IPMK inhibition on mTORC1 signaling as measured by p70S6K mobility shift and S6 phosphorylation (pS6) in INS-1 cells exposed to low glucose (4.5 mM) and stimulated with high glucose (11 mM) for 30 minutes. Also shown are the levels of pAkt, pAMPK, pERK1/2 and mTOR. As positive controls for pAMPK and pAkt, cells were treated with oligomycin (oligo, 5 μM) or stimulated with insulin (100 nM).
**(B)** Mean and standard deviation of pS6 levels from 3 biologically independent samples treated as described in A and normalized by total S6.
**(C)** Mean and standard deviation of the ratio of p70S6K top/bottom bands (phosphorylated /unphosphorylated) from 3 biologically independent samples treated as described in A.
**(D)** Mean and standard deviation of mTOR protein levels from 3 biologically independent samples treated as described in A and normalized by tubulin.
**(E)** Mean and standard deviation of pAkt levels from 3 biologically independent samples treated as described in A and normalized by tubulin.
**(F)** Mean and standard deviation of pAMPK levels from 3 biologically independent samples treated as described in A and normalized by tubulin.
**(G)** Mean and standard deviation of pERK1/2 levels from 3 biologically independent samples treated as described in A and normalized by tubulin (imaging units).

4.5 G indicates basal glucose (4.5 mM) and 11G indicates glucose-stimulated (11 mM); (-) indicates DMSO treated and (+) indicates UNC9750 treated. Data are mean ± SD. Statistical significance is indicated in panels. (*)P=0.01-0.05; (#/*)P=0.07; (#) P>0.07.
